# Optimal CD4T Cell Priming in Lymph Nodes Requires Repertoire Scanning by CD301b^+^ Migratory cDC2 Cells

**DOI:** 10.1101/2020.08.31.276410

**Authors:** Naoya Tatsumi, Alicia L Codrington, Yosuke Kumamoto

## Abstract

Activation of CD4T cells by conventional dendritic cells (cDC) is pivotal in adaptive immunity. However, while the activation mechanism of antigen-specific CD4T cells has been extensively studied, the cellular mechanism that leads to the selection of cognate CD4T cell clones out of the polyclonal pool is incompletely understood. Here, we show that, in the reactive lymph nodes, newly homed naive polyclonal CD4T cells are temporarily retained before leaving the lymph node. This stop-and-go traffic of CD4T cells provides an adequate time window for efficient scanning and timely priming of antigen-specific clones. Mechanistically, upon immunization, CD301b^+^ DCs, a major subset of migratory cDC2 cells, quickly migrate to the draining lymph node and settle in the areas near the high endothelial venules, where they retain incoming polyclonal CD4T cells through MHCII-dependent but antigen-independent mechanisms while concurrently providing cognate stimuli to prime antigen-specific CD4T cells. These results indicate that CD301b^+^ DCs function as an immunological “display window” for CD4T cells to efficiently scan their antigen specificity.

**Graphical Abstract:** 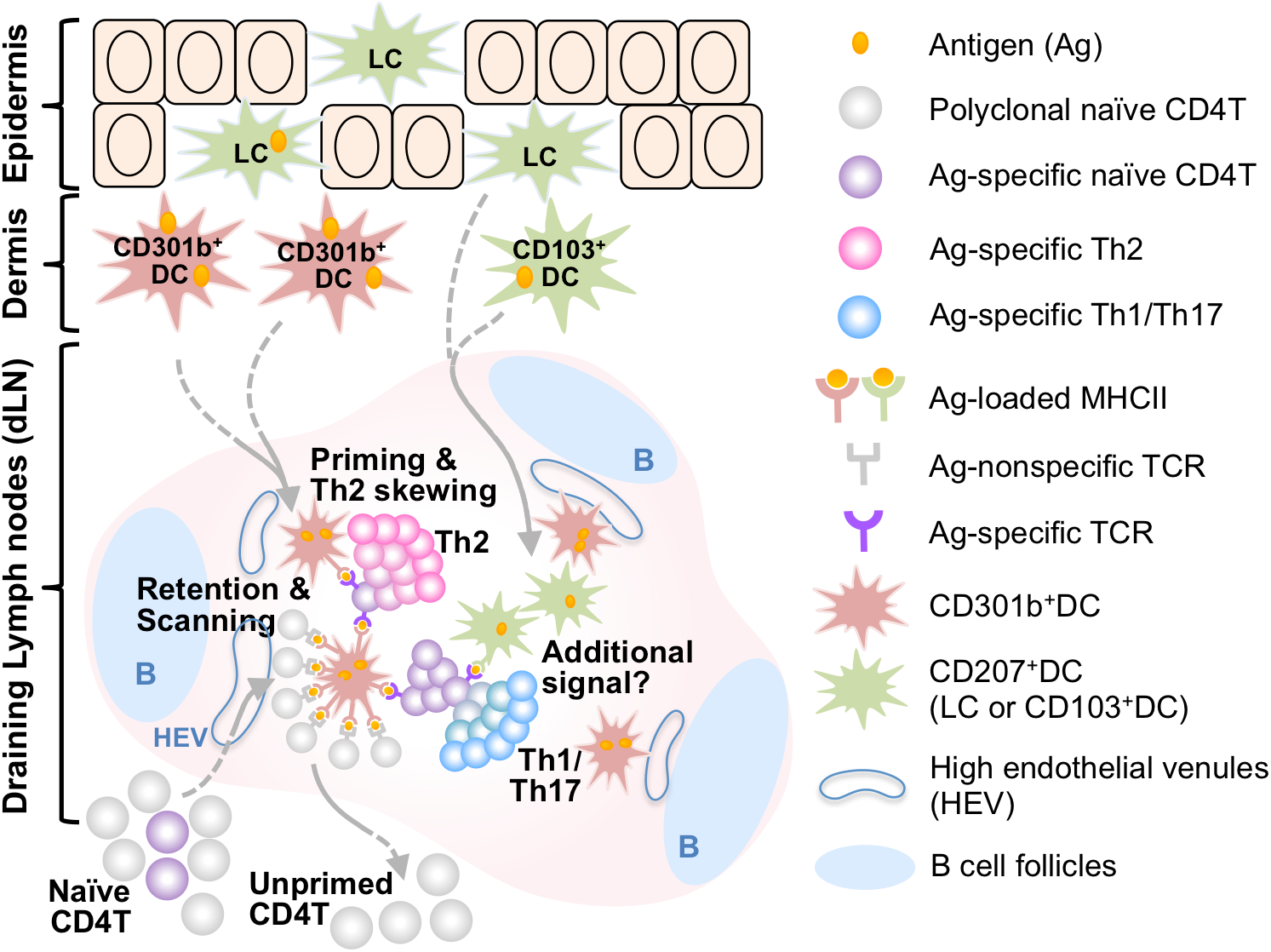

**Highlights:**

- Newly homed polyclonal CD4T cells are temporarily retained in the reactive lymph nodes.
- Depletion of CD301b^+^ DCs results in shorter dwell time of CD4T cells in the draining lymph node and delayed priming of antigen-specific clones.
- The transient retention of polyclonal CD4T cells in the draining lymph node requires MHCII expression on CD301b^+^ DCs but not cognate antigen.
- CD301b^+^ DCs are required for robust expansion of rare antigen-specific CD4T cell clones and their skewing toward Th2 cells.

## Introduction

In the initiation phase of adaptive immune responses, naive T cells undergo a multi-step activation process that results in selective expansion and differentiation of antigen-specific clones. This process consists of homing of polyclonal naive T cells to the draining lymph node (dLN), transient interaction of these polyclonal T cells with antigen-bearing dendritic cells (DCs), engagement of antigen-specific clones with these DCs that leads to a stable interaction, cell cycle entry and rapid proliferation of the antigen-specific T cell clones, and differentiation of the activated T cells in response to cytokines and other antigen non-specific signals from DCs (Bousso, 2008; Catron et al., 2004; Chudnovskiy et al., 2019). Since antigen-specific T cell clones to a given antigen are extremely rare, the scanning of antigen-specificity of polyclonal T cells must be highly efficient in order to meet the demand for combating invading pathogens in a timely manner. Conventional DCs (cDCs) provide the most critical signal to T cells during T cell priming by presenting antigen-derived peptides on their MHC molecules to cognate T cells, but they also provide structural support for maximizing the immune response by inducing LN hypertrophy and promoting the recruitment of polyclonal T cells to the dLN, which collectively leads to higher efficacy in scanning antigen specificity of T cells (Kumamoto et al., 2011; Kumar et al., 2015; Moussion and Girard, 2011).

Naive mouse skin is populated with three major migratory DC subsets, including the epidermal Langerhans cells, dermal CD103^+^ DCs and dermal CD301b^+^ DCs, all of which continuously migrate from the skin to the dLN with subset-specific kinetics and are specialized for the induction of T helper (Th) 17, Th1 and Th2 differentiation of antigen-specific CD4T cells, respectively (Eisenbarth, 2019; Kashem et al., 2017). Although these migratory DCs have long been thought to be critical for initiating T cell responses against skin-borne antigens as they bring antigens from the skin to the dLN, recent studies have challenged this view by demonstrating that dLN-resident DCs can initiate T cell responses before the arrival of the migratory subsets by directly acquiring and presenting antigens that disperse through the lymphatics, especially when the antigen is in a soluble form (Gerner et al., 2017; Gerner et al., 2015; Itano et al., 2003). Thus, the relative contribution of migratory DCs to the initiation of CD4T cell response remains unclear. In addition, while it is well documented that different migratory DC subsets induce different types of Th cell differentiation, their subset-specific roles in the initiation of CD4T cell priming are poorly understood.

CD301b^+^ DCs are a subset of type 2 cDC (cDC2) cells and account for the majority of CD11b^+^ cDCs in the dermis (Gao et al., 2013; Kumamoto et al., 2009; Kumamoto et al., 2013). In hapten-induced contact hypersensitivity, CD301b^+^DCs represent the major hapten-bearing DCs in the dLN 24 hours after sensitization and accumulate at the T-B border areas in the dLN, particularly at areas around the high endothelial venules (HEVs) (Kumamoto et al., 2009). Importantly, transferring hapten-bearing CD301b^+^ DC into naive mice is sufficient for inducing contact hypersensitivity in the recipient animals, suggesting that these DCs are capable of priming naive T cells in the host (Kumamoto et al., 2009; Murakami et al., 2013). More recently, we and others have shown that the differentiation of antigen-specific CD4T cells into Th2 effector cells is selectively impaired when CD301b^+^ DCs were absent or specifically depleted in Mgl2-DTR mice, which express the diphtheria toxin receptor (DTR) under the regulation of the *Mgl2* gene (encoding CD301b) (Gao et al., 2013; Kumamoto et al., 2013; Sokol et al., 2018). However, it was unclear whether the role of CD301b^+^ DC is limited to the induction of Th2 differentiation or they also have a specific role in the CD4T cell priming itself, such as scanning the antigen specificity of polyclonal CD4T cells and triggering cell cycle entry of the cognate clones.

Here, we demonstrate that CD301b^+^ DCs are required for the efficient scanning and timely priming of CD4T cells. In the dLN, CD301b^+^ DCs induce transient retention of polyclonal CD4T cells in the dLN in an MHCII-dependent manner, during which they scan the antigen specificity of CD4T cells and activate cognate clones. When CD301b^+^ DCs were depleted or MHCII expression was abrogated specifically in CD301b^+^ DCs, naive CD4T cells failed to stay in the dLN and the cell cycle entry of antigen-specific clones was significantly delayed. These changes collectively resulted in impaired priming of antigen-specific CD4T cell clones in CD301b^+^ DC-depleted mice, especially when the antigen-specific clones were rare. These results indicate that migratory CD301b^+^ DCs optimize the priming efficacy of CD4T cells.

## RESULTS

### Depletion of CD301b+ DCs Results in Reduced Accumulation of T and B Cells in the dLN

DCs are required for the immunization-induced LN hypertrophy by enhancing the recruitment and/or retention of naïve lymphocytes in the dLN, but subset-specific role of DCs in this process is poorly understood (Kumamoto et al., 2011; Kumar et al., 2015; Moussion and Girard, 2011; Webster et al., 2006). We showed previously that, when mice were depleted of CD301b^+^ DCs and immunized with ovalbumin (OVA) and papain, the size of the dLN early after immunization was smaller than the control animals due largely to the impaired accumulation of CD4T cells, but later it was recovered by the increased germinal center B cells and T follicular helper cells (Kumamoto et al., 2016; Kumamoto et al., 2013). To address if CD301b^+^ DCs directly regulate the recruitment and/or retention of naïve lymphocytes, we transferred CFSE-labeled wild-type (WT) splenocytes retro-orbitally into diphtheria toxin (DT)-treated WT (CD301b^+^ DC-intact) or Mgl2-DTR (CD301b^+^ DC-depleted) mice following subcutaneous immunization with OVA and papain in the right footpad (Figure 1A), which preferentially induces migration of CD301b^+^ DCs to the dLN (Kumamoto et al., 2013). Two hours after the transfer, no significant difference was observed between the WT and Mgl2-DTR recipients in the number of total donor splenocytes, as well as in the numbers of donor CD4T, CD8T, or B cells in both right (draining) and left (non-draining) popliteal LN (Figure 1B), indicating that the depletion of CD301b^+^ DCs has no major impact on the lymphocyte entry into the LNs. Between 2 and 72 hours after the transfer, the numbers of the donor CD4T and CD8T cells in the dLN increased by two-to three-fold in the WT recipients, while the numbers of B cells increased by 30-fold (Figure 1B), suggesting that many T cells that had homed to the dLN left the dLN relatively quickly, while the most of the B cells that had homed to the dLN stayed there throughout this observation period. In the Mgl2-DTR recipients, the numbers of CD4T cells, CD8T cells and B cells were significantly smaller than those in the WT recipients, with the most prominent reduction in CD4T cells (Figure 1B). This was not accounted for by the differences in proliferation of the donor cells, as the majority of the donor cells did not show a significant dilution of the CFSE dye regardless of the CD301b^+^ DC depletion status (Figure 1C), suggesting that the reduction of donor cells in the CD301b^+^ DC-depleted recipients is primarily due to changes in their trafficking rather than proliferation. We previously showed that the DT treatment in Mgl2-DTR mice results in an additional loss of Langerhans cells (which do not express CD301b/*Mgl2*) in the epidermis but not in the skin-dLN due to an unknown mechanism (Kumamoto et al., 2013). However, the reduction of donor cell accumulation is specific to the depletion of CD301b^+^ DCs, as the donor cell numbers were not affected in CD207-DTR mice, in which CD207^+^ DC subsets, including Langerhans cells and dermal CD103^+^ DCs, were depleted (Figure 1D) (Bursch et al., 2007; Ginhoux et al., 2007; Kissenpfennig et al., 2005; Poulin et al., 2007).

**Figure 1.**
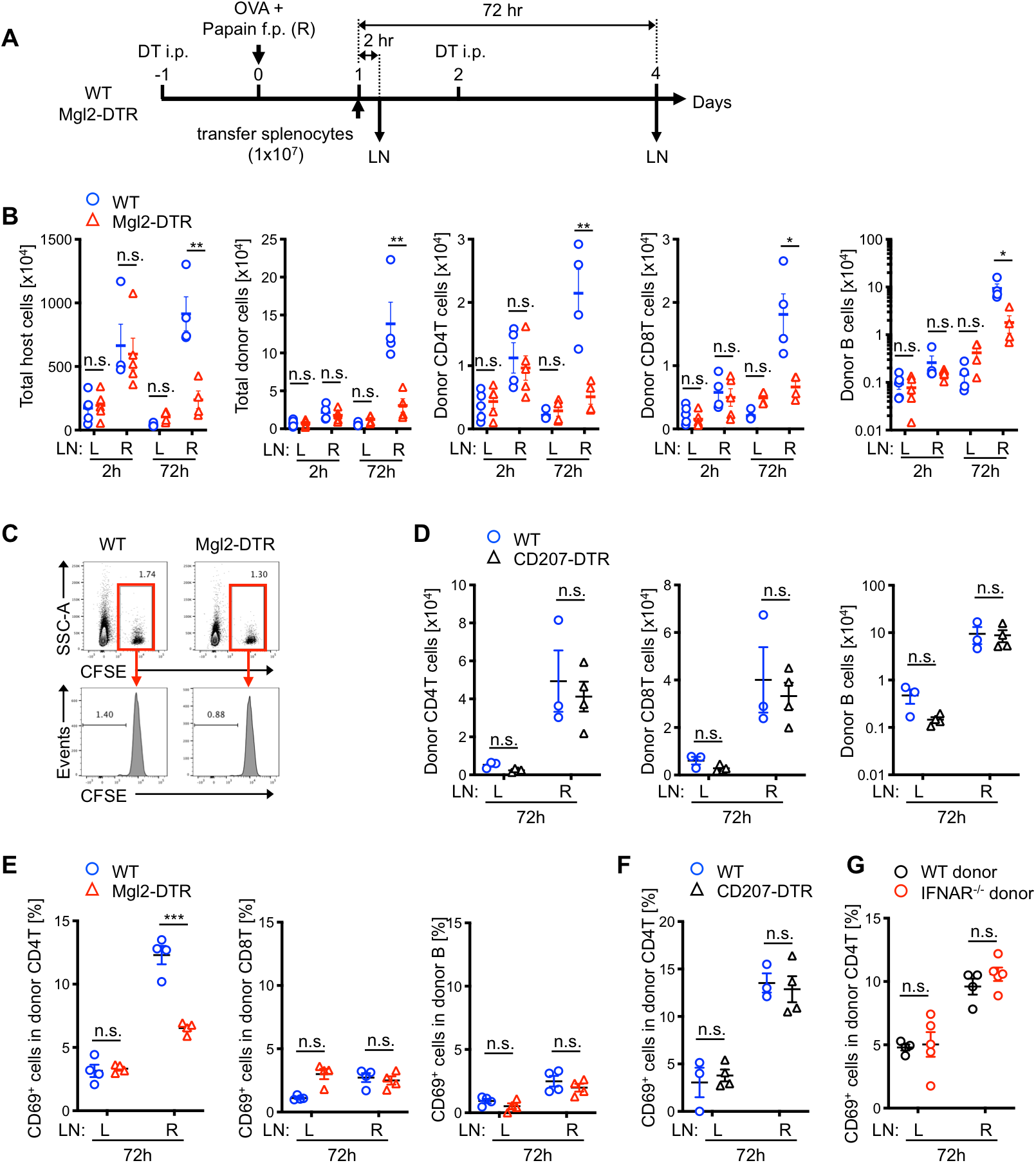
CD301b^+^ DCs are required for lymphocyte accumulation and CD69 upregulation in polyclonal CD4T cells in papain-immunized dLN. (A) Experimental design. CFSE-labeled splenocytes (1 × 10^7^) isolated from naive WT mice were retro-ortbitally transferred into DT-treated WT or Mgl2-DTR mice that had been immunized with OVA and papain in the right hind footpad 24 hours earlier. Right draining (R) and left non-draining (L) popliteal LNs were harvested 2 or 72 hours after the transfer. (B) Number of host- and donor-derived cells recovered from the LNs. (C) CFSE dilution in the donor cells 72 hours after the transfer. (D) As in (A), but the cells were transferred into CD207-DTR mice, which were depleted of Langerhans cells and CD103^+^ DCs. (E-G) Frequency of CD69^+^ cells in the indicated donor cell type 72 hours after the adoptive transfer as in (A). Splenocytes of naive WT mice (E-G) or IFNAR^−/−^ mice (G) were transferred into WT (E-G), Mgl2-DTR (E), or CD207-DTR (F) recipients. Data represent mean ± SEM. n.s., p≥0.05, *p < 0.05, **p < 0.01, ***p < 0.001, by two-tailed Student’s t test.

### CD301b^+^ DCs are required for the upregulation of CD69 in polyclonal CD4T cells recruited to the dLN

Upon entering the inflamed dLN, the newly arrived lymphocytes quickly upregulate CD69, which negatively regulates sphingosine 1 phosphate receptor and inhibits lymphocyte egress from the dLN (Shiow et al., 2006). CD69 is a well-known marker for activated lymphocytes stimulated through their antigen receptors (i.e. TCR or BCR) but is also upregulated in an antigen-nonspecific manner upon activation by inflammatory cytokines such as type I interferons (IFN) (Shiow et al., 2006). We showed previously that immunization with OVA and a Th2-type adjuvant such as papain or alum results in upregulation of CD69 in the endogenous CD4T cells in CD301b^+^ DC-dependent manner (Kumamoto et al., 2013). Consistent with these findings, CD69 was upregulated in the donor CD4T cells in the dLN of WT recipients immunized with OVA and papain, while the upregulation was significantly diminished in the CD301b^+^ DC-depleted recipients (Figure 1E). Notably, among the donor T and B cells, the CD69 upregulation was specific to CD4T cells and was not affected by the depletion of CD207^+^ DCs (Figure 1E, F), suggesting that CD301b^+^ DCs are specifically required for the activation of CD4T cells. The upregulation of CD69 in the donor CD4T cells was independent of type I IFN receptor (IFNAR) in this model, as CD69 was normally upregulated when the donor cells were isolated from IFNAR-deficient mice (Figure 1G), suggesting that the CD69 upregulation in the donor CD4T cells in this model was driven by TCR stimuli or inflammatory cytokines other than type I IFNs.

### CD301b^+^ DCs Retain and Activate Naïve Polyclonal CD4T Cells in the dLN in an MHCII-Dependent Manner

The above data suggest that CD301b^+^ DCs regulate CD4T cell trafficking mainly by facilitating their retention in the dLN. To formally address this possibility, we transferred CFSE-labeled WT CD4T cells into CD301b^+^ DC-intact or depleted mice 24 hours after immunization with OVA and papain in the footpad, and then blocked further LN entry by injecting the anti-CD62L blocking monoclonal antibody (mAb) 2 hours after the donor cell transfer (Figure 2A) (Catron et al., 2006). We first confirmed the blockade of lymphocyte entry into LNs by transferring Cell Tracer Violet (CTV)-labeled CD4T cells immediately after the anti-CD62L mAb injection (Figure S1A). In the spleen, both CFSE-labeled CD4T cells (transferred before the CD62L blockade) and CTV-labeled CD4T cells (transferred after the CD62L blockade) were detected, as homing of naïve T cells to the spleen does not require CD62L (Bradley et al., 1994). In contrast, CFSE-labeled but not CTV-labeled CD4T cells were detected in the LNs, indicating near complete blockade of further CD4T cell entry immediately after the anti-CD62L mAb treatment (Figure S1B). Thus, this experimental setup allowed us to measure the LN dwell time for the donor CD4T cells that had entered the LNs during the initial 2-hour window. In WT recipients, the donor CD4T cells stayed in the dLN for at least 8 hours and began to leave the dLN between 8 and 16 hours after the CD62L blockade (Figure 2B left panel, p=0.45 between 0 and 8 hours, p=0.0011 between 0 and 16 hours). Unexpectedly, while the number of recruited CD4T cell was significantly lower, the kinetics of the donor CD4T cells in the contralateral non-draining (nd) LN followed a similar pattern of that in the dLN, with the majority of donor CD4T cells remaining in the LN for at least 8 hours before leaving (Figure 2B right panel, p=0.6183 between 0 and 8 hours, p=0.0012 between 0 and 16 hours). In contrast, the donor CD4T cells in the CD301b^+^ DC-depleted hosts failed to stay in the dLN and continuously decreased over time, whereas their retention in the ndLN was not impaired by the depletion (Figure 2B). Furthermore, the donor CD4T cells that stayed in the dLN of WT recipients continuously upregulated CD69 during this time, whereas CD69 upregulation in CD301b^+^ DC-depleted hosts was minimal and did not increase between 8 and 16 hours (Figure 2C and 2D). Collectively, these results indicate that CD301b^+^ DCs are required for transient retention and activation of naive polyclonal CD4T cells in the dLN.

**Figure 2.**
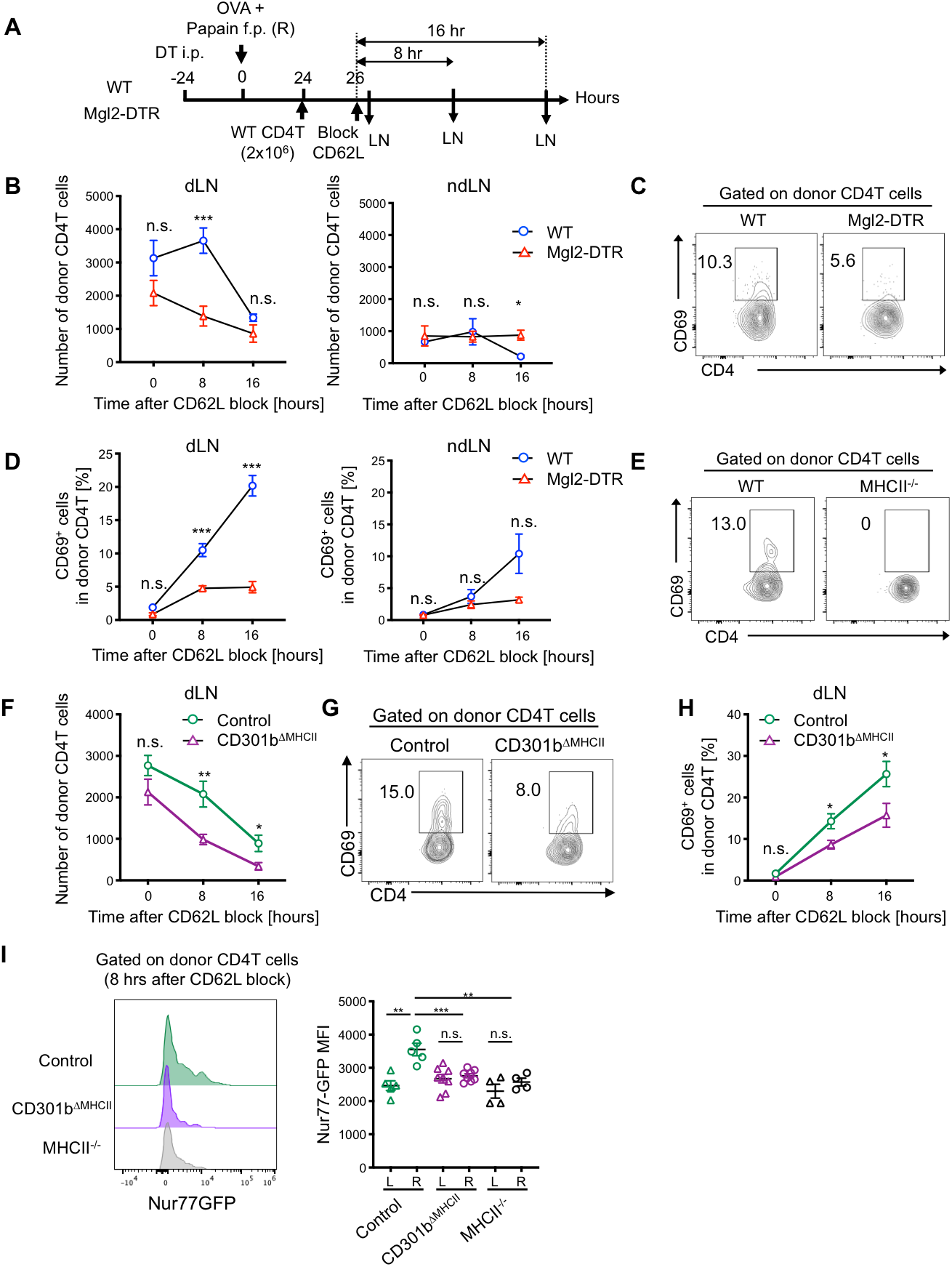
CD301b^+^ DCs Retain and Activate Naive Polyclonal CD4T Cells in the dLN in an MHCII-Dependent Manner. (A) Experimental design. Total CD4T cells were isolated from naive WT donor mice and 2 × 10^6^ cells were transferred into DT-treated Mgl2-DTR or WT mice that had been immunized with OVA and papain in the right footpad 24 hours earlier. Two hours later, further homing of lymphocytes to LNs was blocked by injecting anti-CD62L mAb. The draining and non-draining popliteal LNs were harvested 0, 8 and 16 hours after the CD62L blockade. (B-D) Number (B) and CD69 expression (C, D) of the donor CD4T cells. (E-H) MHCII^−/−^ (*H2ab1*^*−/−*^) mice (E) or *Mgl2*^*+/Cre*^;*H2ab1*^*fl/fl*^ (CD301b^ΔMHCII^) mice (F-H) were transferred with WT CD4T cells as in (A). *Mgl2*^*+/Cre*^;*H2ab1^+/+^* mice were used as control in (F-H). Number (F) and CD69 expression (E, G, H) of the donor CD4T cells are shown. (I) Total CD4T cells isolated from naive Nur77-GFP transgenic mice were transferred into control or CD301b^ΔMHCII^ mice as in (A). The dLNs (R) and ndLNs (L) were harvested 8 hours after the CD62L blockade. Representative FACS histograms (left) and the mean fluorescent intensity (MFI) (right) for the Nur77-GFP reporter expression in the donor CD4T cells are shown. Data represent mean ± SEM (B, D, F, H, I) of 3-9 mice per time-point or representative flow cytometry plot of dLNs 8 hours after the CD62L blockade from at least two independent experiments (C, E, G). *p < 0.05, **p < 0.01, ***p < 0.001, n.s., not significant by two-tailed Student’s t test. See also Figure S1, S2, and S3.

Given the rarity of antigen-specific clones, it seemed unlikely that all CD69^+^ donor CD4T cells (>10% of the donor CD4T cells in the dLN of WT mice 8 hours after the CD62L blockade) were expressing OVA- or papain-specific TCR, but the IFNAR-independent upregulation of CD69 (Figure 1G) nonetheless suggested the involvement of TCR-dependent interaction in this form of CD4T cell activation. Indeed, CD69 upregulation by the donor CD4T cells in the dLN was completely abrogated when the cells were transferred into MHCII-deficient hosts (Figure 2E), further supporting the idea that the interaction between the host MHCII and the TCR on the polyclonal donor CD4T cells is required for the upregulation of CD69 upon dLN entry. Unlike CD8T cells, the intranodal trafficking and LN dwell time of naive CD4T cells are partially governed by the antigen presenting machinery (Fischer et al., 2007; Mandl et al., 2012). In addition, the MHCII-dependent CD69 upregulation in CD4T cells has been shown to correlate with their LN dwell time in naive LNs (Tomura et al., 2010). To directly examine whether MHCII on CD301b^+^ DCs play a role in retaining polyclonal CD4T cells, we generated mice expressing Cre recombinase from the 3’UTR of the *Mgl2* gene (*Mgl2*^*+/Cre*^ mice) and crossed them with mice with the I-Ab alleles flanked by two loxP cassettes (*H2ab1*^*flox/flox*^). The resulting offspring (CD301b^ΔMHCII^ mice) lacks MHCII expression specifically in CD301b^+^ cells (Figure S2A-S2C). Similarly to the CD301b^+^ DC-depleted recipients, the donor polyclonal CD4T cells in CD301b^ΔMHCII^ mice were retained in the dLN less efficiently than those in MHCII-intact hosts (Figure 2F), and the upregulation of CD69 in the donor CD4T cells was also impaired (Figure 2G, 2H and Figure S2D), indicating that MHCII expression in CD301b^+^ DCs plays a significant role in transiently retaining and activating naive polyclonal CD4T cells in the dLN. Similar phenotype was observed when total splenocytes were transferred into CD301b^ΔMHCII^ mice without CD62L blockade (Figure S2D). Notably, the reduction of CD69 upregulation in CD301b^ΔMHCII^ recipients was not as complete as in MHCII-deficient hosts (Figure 2E), suggesting that the remaining MHCII^+^ cells, as well as the residual MHCII expression in CD301b^+^ DCs (Figure S2C), may play a role in CD69 upregulation by CD4T cells in those mice.

Our data thus far indicate the involvement of CD301b^+^ DC-expressed MHCII in retaining naive polyclonal CD4T cells in the dLN and stimulating their CD69 expression, suggesting that CD301b^+^ DCs provide TCR stimulation for CD4T cells upon their homing to the dLN. To further confirm that CD301b^+^ DCs stimulate the TCR of some of those polyclonal CD4T cells, we measured the expression levels of Nur77, a nuclear receptor whose expression levels directly correlate with the TCR signal strength (Au-Yeung et al., 2014; Moran et al., 2011; Zikherman et al., 2012). Since the direct staining of Nur77 with mAb was not sensitive enough to detect its upregulation in the donor polyclonal CD4T cells, we transferred CD4T cells isolated from the Nur77-GFP reporter mice (Moran et al., 2011). In WT recipients, the GFP expression was higher in the dLN than in the contralateral non-draining (nd) LN 8 hours after the CD62L blockade, suggesting that the immunization enhances the TCR stimulation activity in the dLN. In contrast, the GFP expression was similar between the dLN and ndLN in the CD301b^ΔMHCII^ mice as well as in MHCII-deficient mice, indicating that CD301b^+^ DCs are a significant source of the TCR stimulation at this time-point (Figure 2I). Collectively, these results indicate that CD301b^+^ DCs transiently retain and activate polyclonal CD4T cells in the dLN at least partially through MHCII-dependent interactions.

### CD301b^+^ DC-dependent Naive CD4T Cell Retention in the dLN is Irrespective of the CD69 Upregulation or TCR Antigen Specificity

Thus far, our data show that CD301b^+^ DCs are required for the transient retention of naive CD4T cells in the dLNs in mice immunized with a Th2-type adjuvant papain. Our previous studies indicate that CD301b^+^ DC depletion also results in a reduction in CD4T cell accumulation in the dLN in mice immunized with OVA and CpG without making any reduction in CD69 expression (Kumamoto et al., 2013). To examine if CD301b^+^ DCs are required for the retention of polyclonal CD4T cells under non-Th2 immunization conditions, we immunized the mice in the footpad with OVA emulsified in complete Freund’s adjuvant (CFA) (Figure S3A). As observed in mice immunized with papain, the donor CD4T cells in CD301b^+^ DC-depleted mice spent shorter time in the dLNs than those in CD301b^+^ DC-intact mice (Figure S3B). However, CD69 expression levels in the donor CD4T cells were comparable between WT and CD301b^+^ DC-depleted hosts at any time point, suggesting that the differences in CD69 expression levels do not account for the shorter dwell time in this model (Figure S3C).

The above data suggest that, while the interaction between the TCR on CD4T cells and the MHCII in CD301b^+^ DCs is necessary, the cognate interaction involving a specific foreign antigen may not be required for the CD301b^+^ DC-dependent retention of CD4T cells in the dLN. To further clarify the role of cognate and non-cognate interactions in CD301b^+^ DC-dependent CD4T cell trafficking, we co-transferred OVA-specific *Rag1*^*−/−*^ OT-II TCR transgenic CD4T cells and WT naive polyclonal CD4T cells into CD301b^+^ DC-intact or CD301b^+^ DC-depleted mice that had been immunized with OVA and papain for 24 hours and harvested dLN at different time-points to assess their LN dwell time (Figure 3A). This setup allowed us to directly compare the retention of polyclonal and antigen-specific CD4T cells within the same LN. To our surprise, the trafficking kinetics of the antigen-specific CD4T (OT-II) cells was similar to the polyclonal CD4T cells despite the presence of cognate antigen (Figure 3B). In the WT recipients, the numbers of both OT-II cells and WT CD4T cells in the dLN remained similar for at least 8 hours before starting to decrease. In contrast, in the CD301b^+^ DC-depleted recipients, the numbers of both donor cell types continued to decrease, suggesting that CD301b^+^ DCs are required for transiently retaining CD4T cells in the dLN regardless of the presence of cognate antigen.

**Figure 3.**
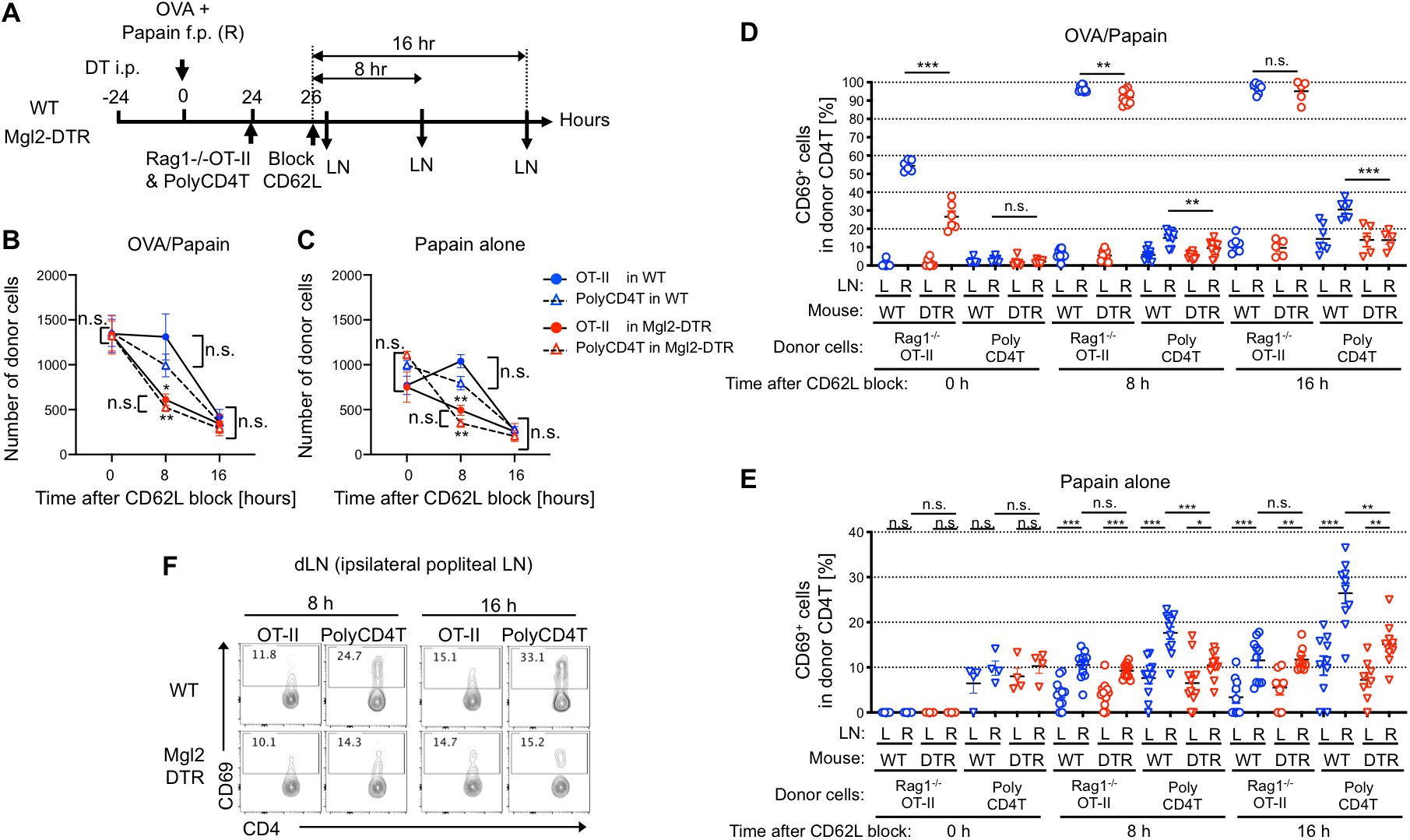
CD301b^+^ DC-dependent Naive CD4T Cell Retention in the dLN is Irrespective of the CD69 Upregulation or TCR Antigen Specificity. (A-C) *Rag1*^*−/−*^ OT-II and WT polyclonal CD4T cells (1 × 10^6^ cells each) were co-transferred into DT-treated Mgl2-DTR or WT recipients that had been immunized with OVA and papain or papain alone 24 hours earlier. The donor cells were allowed to home to LNs for 2 hours, after which further homing was blocked with anti-CD62L mAb. The dLN were harvested 0, 8 and 16 hours after the CD62L blockade (A). Number of donor CD4T cells remained in the dLN of mice immunized with OVA and papain (B) or papain alone (C) at indicated time-points. Pooled data from 4-7 (B) or 3-6 (C) mice per group at each time point are shown. (D-F) *Rag1*^*−/−*^ OT-II and WT polyclonal CD4T cells (1 × 10^6^ cell each) were co-transferred into Mgl2-DTR or WT mice as in (A). Frequency of CD69^+^ cells in the donor CD4T cells in mice immunized with OVA and papain (D) or papain alone (E and F) at indicated time-points. *p < 0.05, **p < 0.01, ***p < 0.001, n.s., not significant by two-tailed Student’s t test. See also Figure S4.

While the results thus far suggest antigen-independency of the CD301b^+^ DC-dependent CD4T cell retention in the dLN, it is still possible that antigen (papain)-specific clones among the donor polyclonal CD4T cells were selectively retained in the dLN in above experiments. To formally exclude this possibility, we repeated the same experiment as in Figure 3A, except that the recipients were immunized with papain alone without OVA so that there was no cognate antigen for the *Rag1*^*−/−*^ OT-II donor cells. Similarly to the results described above, the numbers of both *Rag1*^*−/−*^ OT-II cells and polyclonal CD4T cells retained in the dLN 8 hours after the homing blockade were significantly lower in the CD301b^+^ DC-depleted hosts (Figure 3C). Taken together, these results indicate that CD301b^+^ DCs transiently retain naïve CD4T cells regardless of their TCR specificity.

### CD301b^+^ DC-dependent CD69 Upregulation is Antigen-dependent

Although CD69 is generally thought to promote transient retention of activated T cells in the dLN (Shiow et al., 2006), our experiments using CFA as an adjuvant shows the lack of correlation between the CD301b^+^ DC-dependent CD4T cell retention and the CD69 expression levels (Figure S3). To further clarify the role of antigen recognition in papain-induced CD69 upregulation, we co-transferred *Rag1*^*−/−*^ OT-II and WT polyclonal CD4T cells and compared their CD69 expression levels in mice immunized with either OVA plus papain or papain alone. In mice immunized with OVA plus papain, more than 50% of the OT-II cells in the dLN had already upregulated CD69 at the time of CD62L blockade (2 hours after the OT-II cell transfer) in WT hosts, whereas only about 25% had done so in CD301b^+^ DC-depleted hosts (Figure 3D). Eight hours after the CD62L blockade (10 hours after the transfer), the majority of OT-II cells in the dLN expressed CD69 in both WT and CD301b^+^ DC-depleted recipients, but its expression levels were slightly but significantly reduced in the latter (Figure 3D, see also Figure 5H). Given that many OT-II cells fail to stay in the dLN of CD301b^+^ DC-depleted recipients during this time, the data suggest that the CD69 expression alone is not sufficient to prevent CD4T cells from leaving the dLN of those mice. As shown in Figure 2, CD69 upregulation was also observed in the donor polyclonal CD4T cells in the dLN of WT recipients, but was significantly impaired in the CD301b^+^ DC-depleted recipients (Figure 3D). Similarly, the CD69 upregulation in the donor polyclonal CD4T cells was CD301b^+^ DC-dependent in mice immunized with papain alone (Figure 3E and 3F). However, even though there was a clear, antigen-nonspecific upregulation of CD69 induced by papain alone in the donor *Rag1*^*−/−*^ OT-II cells in the dLN compared to those in the ndLN, it was not impaired by the depletion of CD301b^+^ DCs (Figure 3E and 3F). The antigen-dependent and independent CD69 upregulation was also observed in *Rag1^+/−^* OT-II cells when *Rag1*^*−/−*^ OT-II, *Rag1^+/−^* OT-II and polyclonal CD4T cells were co-transferred (Figure S4). Taken together, these results indicate that, upon immunization with papain, CD69 is upregulated in CD4T cells in both antigen-dependent and independent manner. While the requirement of CD301b^+^ DCs for the CD4T cell retention in the dLN is independent of the TCR specificity (Figure 3C), CD301b^+^ DCs are required only for the antigen-dependent CD69 upregulation in CD4T cells (Figure 3D-3F), again suggesting that the impaired CD69 upregulation alone does not account for the shorter dLN dwell time of CD4T cells in CD301b^+^ DC-depleted mice. Since *Rag1*^*−/−*^ and *Rag1^+/−^* OT-II cells behaved similarly, we used *Rag1^+/−^* OT-II cells to examine the role of CD301b^+^ DCs in antigen-specific CD4T cell responses in the following experiments.

### CD301b^+^DCs Directly Present Soluble Foreign Antigens to CD4T cells in the dLN Immediately After Their Homing

The above data suggest that CD301b^+^DCs provide CD4T cells with (1) MHCII-dependent but antigen-independent signal to transiently stay in the dLN and (2) MHCII- and antigen-dependent signal to upregulate CD69 expression. This latter signal likely represents the direct antigen presentation to the cognate CD4T cell clones. We and others showed previously that, in a model of hapten-induced contact hypersensitivity, CD301b^+^ DCs are enriched at the T-B cell border area in the dLN and particularly at areas surrounding HEVs, whereas CD207^+^ DCs (including epidermal Langerhans cells and dermal CD103^+^ DCs) are segregated from the CD301b^+^ DCs and localized more preferentially to the deeper T cell zone (Kissenpfennig et al., 2005; Kumamoto et al., 2009; Stoltzfus et al., 2020), suggesting that CD301b^+^ DCs are the first DC population encountered by CD4T cells as they home to the LN through the HEVs. In the dLNs of WT mice immunized with OVA and papain, CD301b^+^ DCs were also enriched at the T-B boundary areas and located in closer proximity the HEVs than CD207^+^ DCs (Figure 4A-4C). When mice were immunized with fluorescently labeled OVA and papain, CD301b^+^ DCs were the most efficient in taking up the OVA among the antigen presenting cells in the dLN during the first 3 days after immunization (Figure 4D, S5A and S5B). Although OVA^+^ B cells outnumbered any OVA^+^ DC subset due to the abundance of the total B cells, the amount of OVA per cell in B cells was minimal, suggesting that B cells are not the major antigen-presenting cells during these time-points (Figure 4E and S5B). Notably, when the mice were immunized with OVA and CFA, both CD301b^+^ DCs and CD103^+^ DCs took up the antigen at similar levels (Figure S5C).

**Figure 4.**
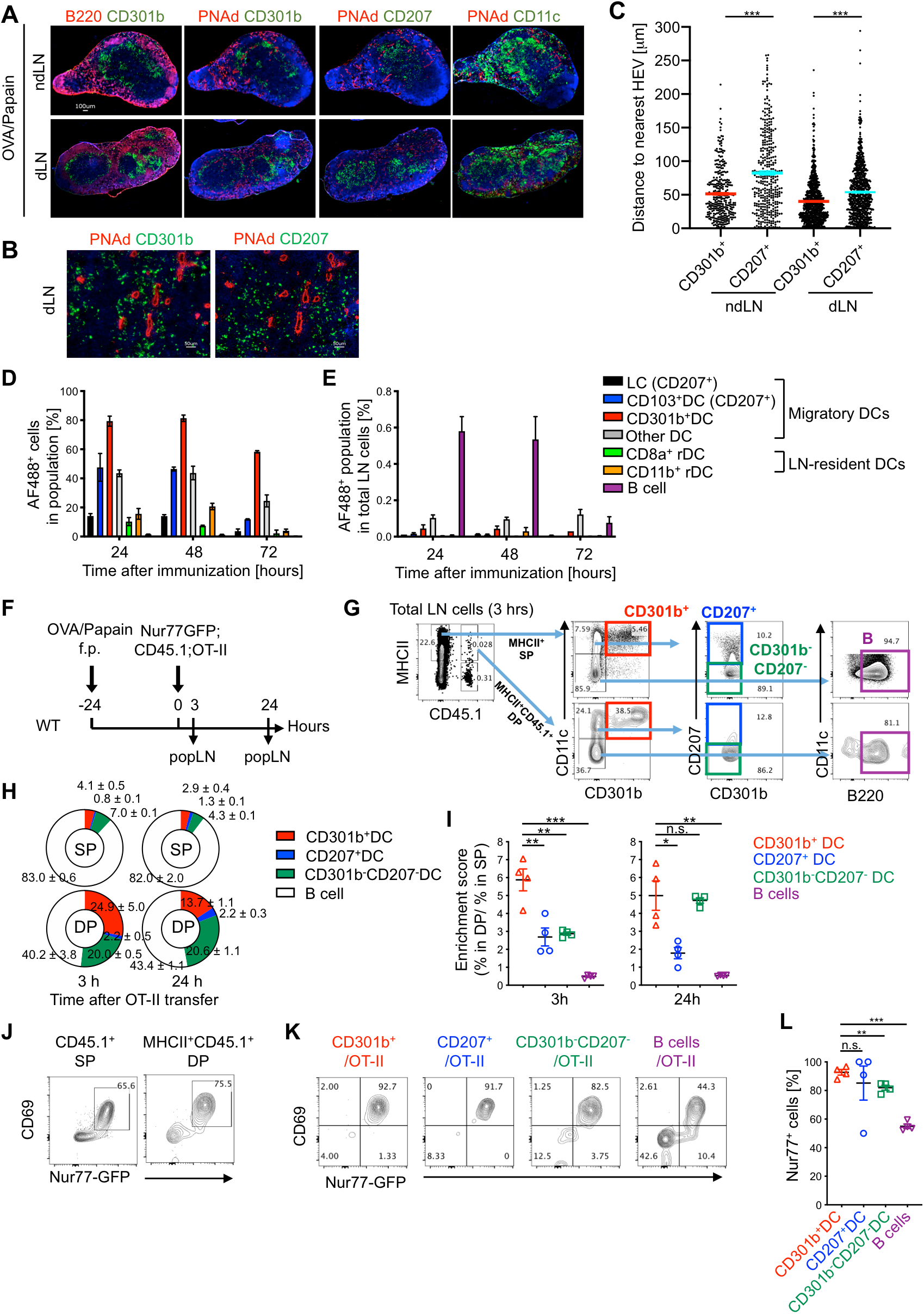
CD301b^+^ DCs Directly Present Soluble Foreign Antigens to CD4T cells in the dLN Immediately After Their Homing. (A-C) WT mice were immunized with OVA and papain in the footpad. Twenty-four hours later, the dLN and contralateral ndLN were harvested and stained for DC subsets (CD301b, CD207, CD11c), HEV (PNAd) and B cell follicles (B220). Representative images (A, B) and the distance of CD301b^+^ or CD207^+^ cells to the nearest HEV (C) are shown. (D, E) Mice were immunized with Alexa Fluor 488 (AF488)-labeled OVA protein with papain in the footpad and the dLNs were harvested at indicated time-points. Frequency of AF488^+^ cells among each cell subset (D) and frequency of each AF488^+^ population among total LN cells (E) are shown. LC, Langerhans cell; rDC, LN-resident DC. n=3-4 per each time-point. Data are pooled from two independent experiments and represented as mean ± SEM. (F-L) Detection of DC-T cell conjugates *ex vivo*. Nur77-GFP;CD45.1;OT-II cells (5 ×10^6^ cells) were transferred into WT mice 24 hours after immunization with OVA and papain. The dLNs were harvested 3 and 24 hours after the transfer and analyzed for the host (CD45.1^−^) MHCII^+^ cell-donor OT-II cell (CD45.1^+^) conjugates by flow cytometry as in (F). Flow cytometric events were analyzed according to the gating strategy for DC and B cell subsets shown in (G). (H) Composition of the MHCII^+^ cells (CD301b^+^ DCs, CD207^+^ DCs, CD301b^−^ CD207^−^ DCs, and B220^+^ CD11c^−^ B cells) within the CD45.1^−^ MHCII^+^ single-positive (SP) or CD45.1^+^ MHCII^+^ double-positive (DP) flow cytometric events in the dLN 3 (left) and 24 hours (right) after the OT-II cell transfer into mice immunized with OVA and papain. (I) Enrichment of CD301b^+^ DCs within the host MHCII^+^ cell-donor OT-II cell conjugates. Enrichment scores were calculated as [% in CD45.1^+^ MHCII^+^ DP / % in MHCII^+^ SP] for each cell type. (J) Expression of CD69 and Nur77-GFP reporter in the CD45.1^+^ MHCII^−^ SP or CD45.1^+^ MHCII^+^ DP events in the dLNs harvested 3 hours after the OT-II cell transfer. (K, L) Expression of the Nur77-GFP reporter in the OT-II cells conjugated with indicated cell type in the dLN 3 hours after the OT-II cell transfer. Data indicate mean ± SEM (D, E, I, L) or representative flow cytometry plots of at least two independent experiments (G, J, K). *p < 0.05, **p < 0.01, ***p < 0.001, n.s., not significant by two-tailed Student’s t test. See also Figures S5 and S6.

Next, to examine whether CD301b^+^ DCs directly interact with the antigen-specific CD4T cells in the dLN, we attempted to detect the cellular conjugates between CD4T cells and CD301b^+^ DCs formed in the dLN by flow cytometry. To that end, we transferred Nur77-GFP;CD45.1;OT-II cells into WT (CD45.1^−^) recipients immunized with OVA and papain one day prior, and then harvested and analyzed all dLN cells 3 and 24 hours after the transfer without excluding doublets (Figure 4F). A small fraction of CD45.1^+^ MHCII^+^ doublet events was detected, indicative of the stable interactions between the CD45.1^+^ OT-II cells and the MHCII-expressing host cells (Figure 4G and S6A). The majority of these doublets were formed *in vivo* in the dLN and not *ex vivo* in the cell suspension, as no fluorochrome exchange was observed when two sets of LN cells were separately stained for CD45.1 and MHCII with a different set of fluorochromes and mixed together *in vitro* as previously described (Figure S6B) (Giladi et al., 2020; Reinhardt et al., 2009). The CD45.1^+^ MHCII^+^ doublets were significantly reduced when the mice were immunized with papain alone, indicating the antigen-dependency of this interaction (Figure S6C-S6E). Although a significant portion of the CD45.1^+^ MHCII^+^ doublets was contributed by B220^+^ CD11c^−^ B cells, this interaction seems to be antigen-independent, since similar interaction was detectable after immunization with papain alone, whereas the DC-OT-II cell interaction was largely dependent on the presence of OVA (Figure S6C-S6E). Three hours after the OT-II cell transfer, approximately 25% of the CD45.1^+^ MHCII^+^ doublets contained CD301b^+^ DCs (Figure 4H). When compared with the DC subset composition in the singlet MHCII^+^ cells (i.e., MHCII^+^ cells not conjugated with OT-II cells), CD301b^+^ DCs were significantly more enriched in the OT-II doublets over CD207^+^ and double negative (CD301b^−^ CD207^−^) DCs or B cells (Figure 4I). While the enrichment of CD301b^+^ DCs in the CD45.1^+^ MHCII^+^ doublets was similar between 3 and 24 hours after the OT-II cell transfer, there was an increase in the enrichment of CD301b^−^ CD207^−^ DCs for the latter time-point, suggesting that the donor OT-II cells primarily interact with CD301b^+^ DCs early after their homing to the dLN, but later they gain interaction with other DC subsets (Figure 4H and 4I). Notably, the majority of OT-II cells conjugated with any DC subset 3 hours after the transfer expressed the Nur77-GFP reporter, whereas only about a half of the OT-II cells conjugated with B cells expressed this reporter (Figure 4J-4L), indicating that DCs but not B cells are the primary antigen presenting cells at this time-point. These results suggest that CD301b^+^ DCs directly present soluble foreign antigens to cognate CD4T cells immediately after their homing to the dLN.

### CD301b^+^ DCs are Required for Timely Priming of Antigen-Specific CD4T Cells

To examine the role of CD301b^+^ DCs in antigen-specific CD4T cell priming and the subsequent expansion, we primed CFSE-labeled OT-II cells in WT or CD301b^+^ DC-depleted mice and monitored their activation kinetics. The priming was semi-synchronized by blocking CD62L two hours after transferring OT-II cells into mice immunized with OVA and papain one day prior (Figure 5A). In both CD301b^+^ DC-intact and depleted mice, the majority of OT-II cells remained undivided for up to 16 hours post CD62L blockade (Figure 5B). More than a half of the OT-II cells underwent the first cell division between 16 and 32 hours post CD62L blockade in WT mice (Figure 5B and 5C). In CD301b^+^ DC-depleted mice, however, only about 20% of the OT-II cells have divided during this period, suggesting a significant delay in their initial cell cycle entry (Figure 5B and 5C). This was also reflected in the significantly fewer cell division cycles at 56 hours post CD62L blockade (Figure 5B and 5C). As was observed in mice co-transferred with OT-II and WT CD4T cells (Figure 3), the number of OT-II cells remained in the dLN 8 hours after the CD62L blockade was significantly lower in the CD301b^+^ DC-depleted mice than in WT mice, but the difference between the two groups disappeared as the majority of the OT-II cells left the dLN by 16 hours after the homing blockade (Figure 5D inset). However, likely due to the delayed cell division, there was only a minimal expansion of OT-II cells in CD301b^+^ DC-depleted mice by the 56 hour time-point, while the number of OT-II cells was dramatically increased in WT mice between 32 and 56 hours post CD62L blockade (Figures 5D). This also resulted in a significant reduction in the percentage of OT-II cells within the total CD4 T cell population in CD301b^+^ DC-depleted mice (Figure 5E). The reduction of OT-II cells was also partially accounted for by the increased cell death in OT-II cells (Figure 5F). These results collectively indicate that CD301b^+^ DCs are required for the timely priming and maximal expansion of antigen-specific CD4T cells during the early priming phase.

**Figure 5.**
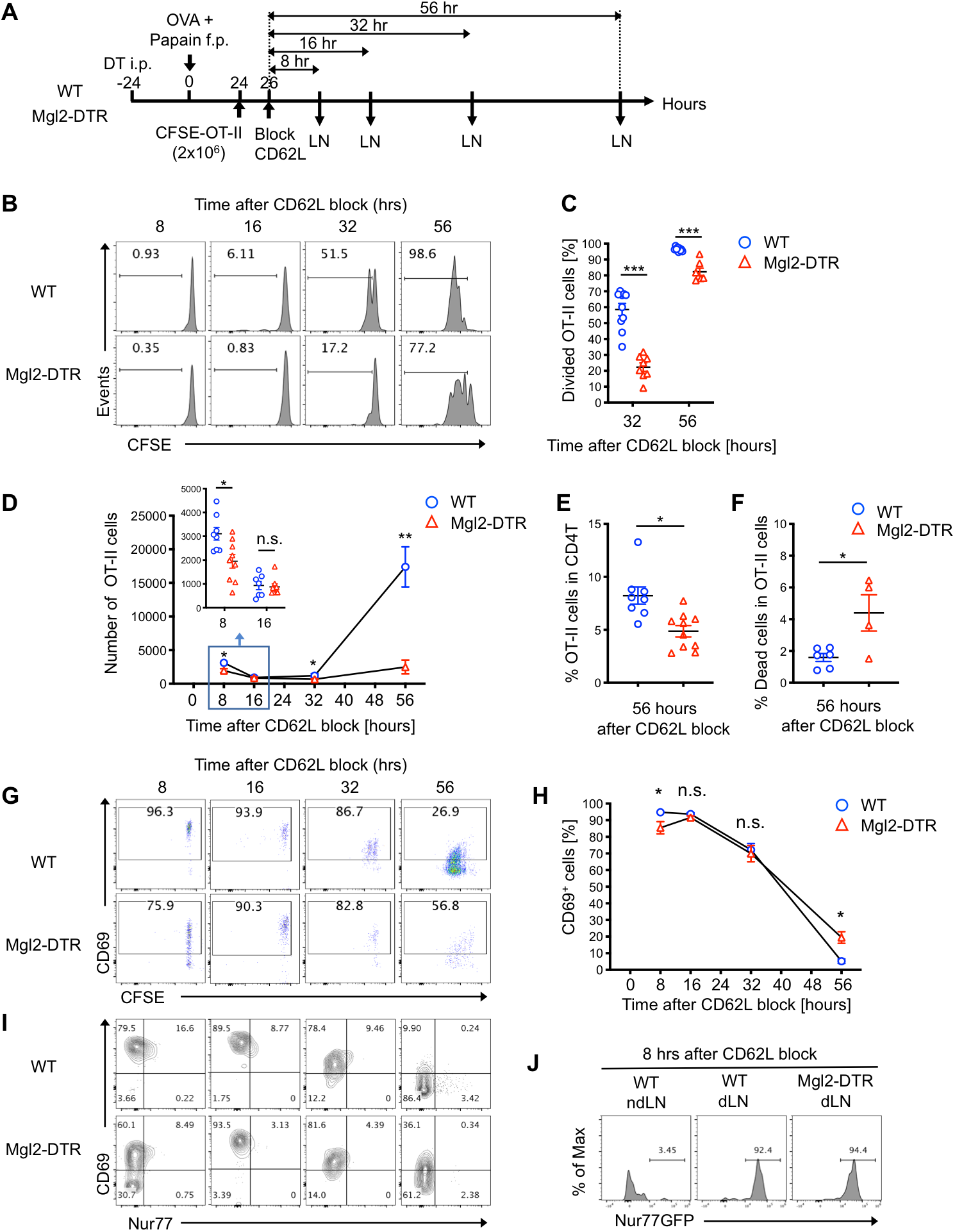
CD301b^+^ DCs are Required for Timely Priming of Antigen-Specific CD4T Cells. (A-I) CFSE-labeled OT-II cells (2 × 10^6^ cells) were transferred into DT-treated Mgl2-DTR or WT mice 24 hours after immunization and allowed to home to LNs for 2 hours, after which further homing was blocked with anti-CD62L mAb. dLNs were harvested at indicated time-points after the CD62L blockade as in (A). Cell division (B, C), cell numbers (D), percentage among total CD4T cells (E), cell death (F), CD69 expression (G, H) and Nur77 expression (I) in the donor OT-II cells at indicated time-points are shown. The inset in (D) shows the number of OT-II cells in the dLN 8 and 16 hours after the CD62L blockade. (J) Nur77-GFP;CD45.1;OT-II cells (2 × 10^6^ cells) were transferred into recipient mice as in (A) and the dLNs were harvested 8 hours after the CD62L blockade. Nur77-GFP reporter expression of the donor OT-II cells is shown. Data represent mean ± SEM (C, D, E, F, H) or show representative flow cytometry plots of at least two independent experiments (B, G, I, J). In (D) and (H). Data are pooled from 4–9 mice per group at each time-point. *p < 0.05, **p < 0.01, ***p < 0.001, n.s. not significant by two-tailed Student’s t test. See also Figure S7

To gain molecular insights, we next analyzed the expression of early activation markers in OT-II cells during the early priming phase. Eight hours after the CD62L blockade, more than 90% of the OT-II cells in WT recipients had already upregulated CD69, and a part of the CD69^+^ OT-II cells at this time-point co-expressed Nur77 (Figure 5G-5I). The expression of CD69 and Nur77 gradually decreased over time and returned to the basal level by 56 hours post CD62L blockade. As in Figure 3D, the CD69 expression in the donor OT-II cells in CD301b^+^ DC-depleted mice was significantly reduced at earlier time-points but was later recovered, suggesting a delay but not loss of TCR stimulation (Figure 5G and 5H). In contrast, the Nur77 expression in the donor OT-II cells remained significantly lower in the CD301b^+^ DC-depleted recipients than in WT recipients throughout the time-points analyzed, suggesting attenuation in the TCR signal strength (Figure 5I). Similarly, priming of OT-II cells in the CD301b^ΔMHCII^ mice also resulted in a significant delay in cell cycle entry and CD69 upregulation as well as in impaired Nur77 expression (Figure S7). However, the use of Nur77-GFP;OT-II cells did not further clarify these differences due to its high sensitivity, as most of the OT-II cells expressed GFP at the brightest levels regardless of the CD301b^+^ DC depletion, which further supports our interpretation that TCR stimulation is not completely absent in the CD301b^+^ DC-depleted recipients (Figure 5J). Taken together, these results indicate that CD301b^+^ DCs are required for the timely and maximal stimulation of antigen-specific CD4T cell clones, but other antigen presenting cells can eventually provide TCR stimulation even if CD301b^+^ DCs are depleted.

### Early Interaction with CD301b^+^ DCs is Critical for the Maximal Expansion and the Fate Decision by Antigen-specific CD4T Cells

Since the data thus far indicated the crucial role of CD301b^+^ DCs in the initial activation and cell cycle entry of antigen-specific CD4T cells, we hypothesized that there is a critical time-window in which CD301b^+^ DCs are required. To test this possibility, we depleted CD301b^+^ DCs at two different time-points, before or after the initial cell division, and then analyzed cell division cycles of the transferred OT-II cells at 56 hours after the CD62L blockade (Figure 6A). Similarly to the above experiments in which CD301b^+^ DCs were depleted one day *before* the immunization (Figure 5A), the depletion of CD301b^+^ DCs one day *after* the immunization (at the same time with the OT-II cell transfer) resulted in fewer cell divisions when analyzed 56 hours after their dLN entry (Figure 6B). In contrast, there was no difference in OT-II cell divisions between WT and CD301b^+^ DC-depleted mice when CD301b^+^ DC were depleted two days *after* the immunization (Figure 6B), which is roughly the same time as many OT-II cells undergo initial cell divisions in WT mice (Figure 5B). These results indicate that CD301b^+^ DCs are required for the timely cell cycle entry of antigen-specific CD4T cells rather than their continued proliferation.

**Figure 6.**
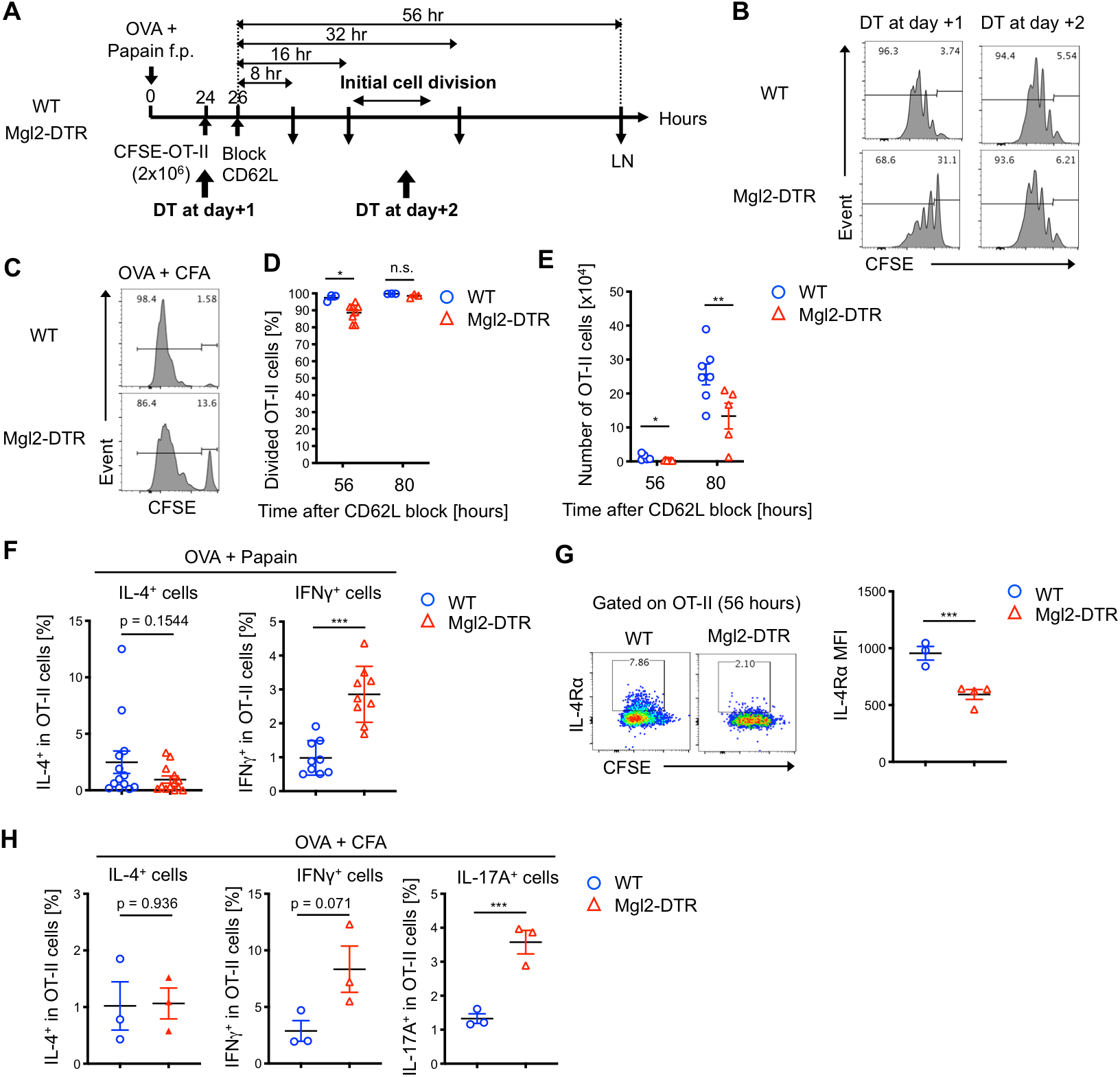
Early Interaction with CD301b^+^ DCs is Critical for the Maximal Expansion and the Fate Decision by Antigen-specific CD4T Cells. (A,B) CFSE-labeled OT-II cells (2× 10^6^ cells) were transferred into Mgl2-DTR or WT mice 24 hours after immunization with OVA and papain and allowed to home to LNs for 2 hours, after which further homing was blocked with anti-CD62L mAb. The recipient mice were treated with DT 1 or 2 days after the immunization and the dLNs were harvested 56 h after the CD62L blockade as in (A). Cell division profile of the OT-II cells is shown in (B). (C-E) CFSE-labeled OT-II cells (2 × 10^6^ cells) were transferred into DT-treated Mgl2-DTR or WT mice 24 hours after immunization with OVA and CFA and allowed to home to LNs for 2 hours, after which further homing was blocked with anti-CD62L mAb. The dLNs were harvested 56 or 80 hours after the CD62L blockade. Cell division (C, D), cell number (E) of the OT-II cells in the dLN at indicated time-points are shown. Histograms in (C) show representative data at the 56-hour time-point. (F-H) OT-II cells (2 × 10^6^ cells) were primed with OVA and papain as in Figure 5A (F, G) or with OVA and CFA as in Figure 6C (H). dLN cells were harvested 80 hours after the CD62L blockade and stimulated *ex vivo* with PMA and ionomycin for intracellular cytokine staining (F,H). Alternatively, the cell surface expression of IL-4Rα in OT-II cells was examined by flow cytometry 56 hours after the CD62L blockade (G). Frequencies of IL-4^+^, IFNγ^+^, IL-17A^+^ or IL-4Rα^+^ cells among the donor OT-II cells are shown. Data represent mean ± SEM (D-H) or show representative flow cytometry plot of at least two independent experiments (B, C, G). *p < 0.05, **p < 0.01, ***p < 0.001, n.s. not significant by two-sided Student’s t test. See also Figure S8

To examine whether the requirement of CD301b^+^ DCs for the optimal expansion of antigen-specific CD4T cells is specific to Th2 conditions, we next immunized the mice with OVA and CFA. Similarly to the mice immunized with OVA and papain, OT-II cells in the CD301b^+^ DC-depleted mice showed a small but significant reduction in the number of cell divisions 56 hours after the CD62L blockade compared to those in the WT mice (Figure 6C and 6D), which also lead to a reduction in the number of OT-II cells in the dLN (Figures 6E). These results indicate that CD301b^+^ DCs are required for the optimal priming and maximal expansion of antigen-specific CD4T cells under both Th2 and non-Th2 immunization conditions.

We previously reported that the depletion of CD301b^+^ DCs abolishes Th2 cell differentiation of antigen-specific CD4T cells, while their differentiation into Th1 cells remains largely unaffected (Kumamoto et al., 2013). The requirement of CD301b^+^ DCs for Th2 differentiation is specific to this DC subset, as the depletion of CD207^+^ DCs (including epidermal Langerhans cells and dermal CD103^+^ DCs) did not affect Th2 differentiation (Figure S8). However, given that the critical time-window for the interaction between CD301b^+^ DCs and antigen-specific CD4T cells is narrow (Figure 6B), it is possible that the lagged priming of recirculating OT-II cells had masked the impact of CD301b^+^ DC depletion on Th1 differentiation in our previous experiments, as those experiments did not employ the semi-synchronized priming model used in the above experiments. To examine the direct impact of the CD301b^+^ DC depletion on the fate decision by antigen-specific CD4T cell clones primed in a semi-synchronized setting, we transferred OT-II cells and immunized mice as in Figure 5A. The dLNs were harvested 80 hours after the CD62L blockade and re-stimulated *ex vivo* with PMA and ionomycin. While the reduction in IL-4^+^ Th2 cells did not reach statistical significance at this time-point, their differentiation into IFNγ^+^ Th1 cells was significantly enhanced in CD301b^+^ DC-depleted mice (Figure 6F). In addition, the expression of IL-4 receptor α in the OT-II cells was significantly reduced in the CD301b^+^ DC-depleted mice 56 hours after the CD62L blockade, suggesting that the OT-II cells had already been skewed against Th2 differentiation in the CD301b^+^ DC-depleted mice by that time (Figure 6G). Similarly, when mice were immunized with OVA and CFA, the expression of IFNγ and IL-17A in OT-II cells was enhanced by the depletion of CD301b^+^ DCs (Figure 6H). These results collectively suggest that early interaction with CD301b^+^ DCs during the priming of antigen-specific CD4T cells generally promotes Th2 differentiation while suppressing their differentiation into Th1 and Th17 cells.

### CD301b^+^ DCs are Required for Optimal Priming and Expansion of Rare Antigen-specific CD4T Cell Clones

Under physiological conditions, DCs perform a daunting task, scanning millions of naive CD4T cells to find a few antigen-specific clones. Since our data thus far indicated that CD301b^+^ DCs are required for the transient retention of polyclonal CD4T cells in the dLN and timely priming and expansion of antigen-specific clones, we hypothesized that their role is even more critical when the antigen-specific CD4T cell clones are rare. To test this hypothesis, we transferred titrated numbers of OVA-specific OT-II cells (1, 10, 100, 1000, or 10,000 cells) into CD301b^+^ DC-intact or depleted mice that were immunized with OVA and papain for 24 hours without CD62L blockade and enumerated OT-II cells in the dLN 4 days after the transfer (Figure 7A). The numbers of recovered OT-II cells were constantly smaller in CD301b^+^ DC-depleted mice than in CD301b^+^ DC-intact mice, which was more exaggerated when the input OT-II cell numbers were lower (100 or 1000 input cells compared to 10,000 input cells, Figure 7B and 7C). Similarly, the numbers of recovered OT-II cells were constantly smaller in CD301b^ΔMHCII^ mice compared to the MHCII-intact recipients when a lower number of OT-II cells were transferred (Figure 7D), indicating that CD301b^+^ DCs maximize the priming efficiency of rare antigen-specific CD4T cell clones through MHCII-dependent interactions. The potential controversy between the results described in Figure 5D (that the CD301b^+^ DC depletion results in impaired expansion of OT-II cells) and the results from the mice with higher input OT-II cell numbers (showing no significant reduction in the output OT-II cell numbers when 10,000 OT-II cells were transferred) is likely due to the lack of CD62L blockade in the latter, as lagged priming of the recirculating OT-II cells can mask the impaired priming efficacy especially when the number of recirculating OT-II cells is high.

**Figure 7.**
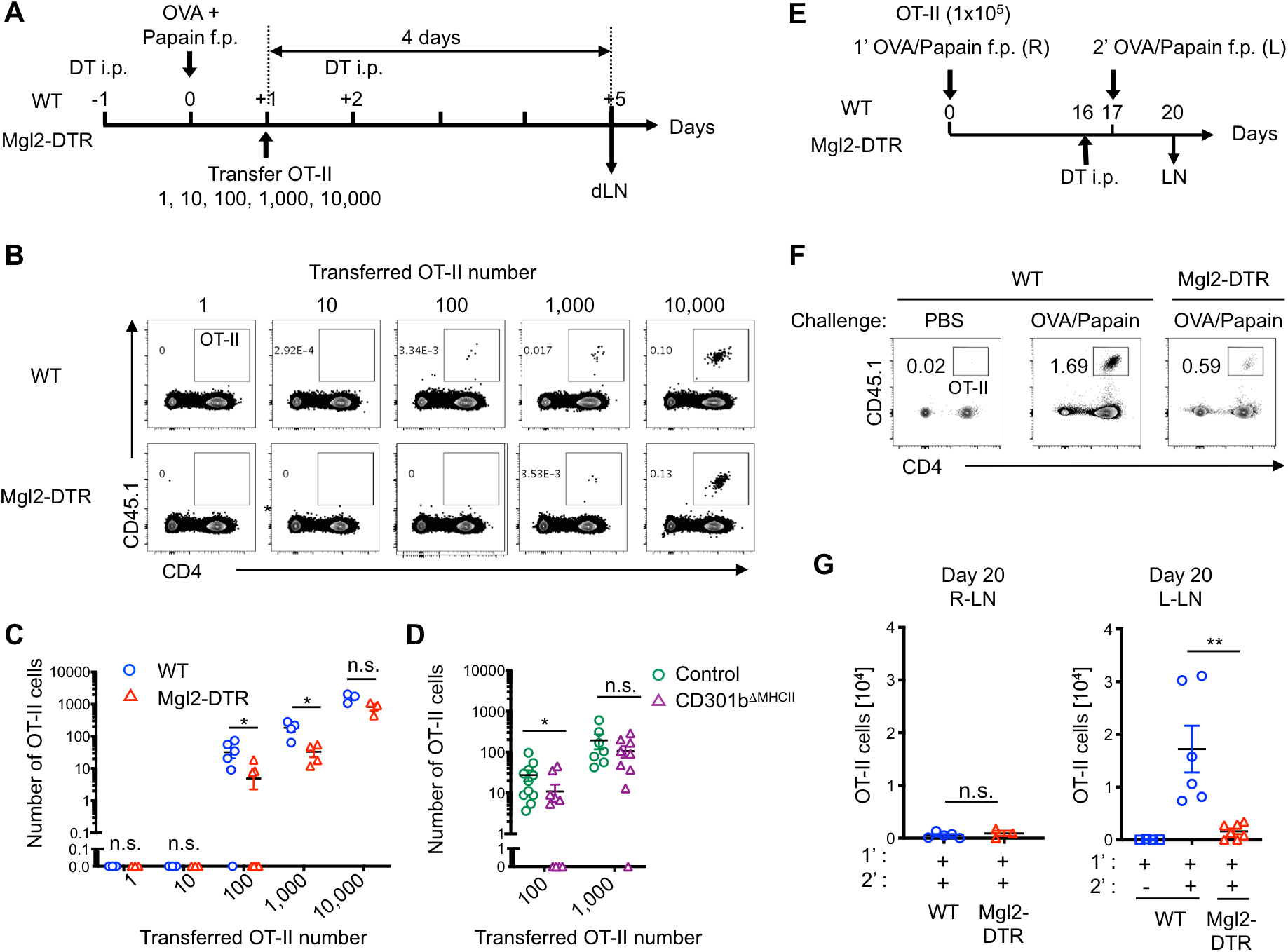
CD301b^+^ DCs are Required for Optimal Priming and Expansion of Rare Antigen-specific CD4T Cell Clones. (A-C) Titrated numbers (1, 10, 100, 1,000, or 10,000) of CFSE-labeled OT-II cells together with unlabeled WT CD4T cells (to match the total donor cell number always to 10,000) were transferred into DT-treated Mgl2DTR or WT mice 24 hours after immunization with OVA and papain, and the dLNs were harvested 4 days after the adoptive transfer (A). Flow cytometric plots of the total LN cells (B) and the number of OT-II cells detected in the dLN (C) are shown. Data are pooled from 3–6 mice per group. (D) CFSE-labeled naive OT-II cells (100, or 1,000) together with unlabeled WT CD4T cells were transferred into control or CD301b^ΔMHCII^ mice 24 hours after immunization with OVA and papain. The number of OT-II cells detected in the dLNs 4 days after the immunization is shown. Data pooled from 7–11 mice per group. (E-G) WT or Mgl2-DTR mice were transferred with 1 × 10^5^ OT-II cells and immunized with OVA and papain in the right footpad on day 0. The mice were treated with DT on day 16 and re-challenged with OVA and papain in the left footpad on day 17. Popliteal LNs were harvested on day 20, 3 days after the secondary challenge (E). Flow cytometry plots of the total dLN cells in the left popliteal LNs (F) and the number of OT-II cells detected in the right (R) or left (L) popliteal LNs are shown. (B, F) Data are representative. Data represent means ± SEM (C, D, G) or show representative flow cytometry plots of at least two independent experiments (B, F). *p < 0.05, n.s., not significant,by two-tailed Student’s t test. See also Figure S9.

To further confirm the role of CD301b^+^ DCs in maximizing the priming efficacy of rare antigen-specific CD4T cell clones, we transferred a high number (1×10^5^) of OT-II cells into WT or Mgl2-DTR recipients, immunized with OVA and papain in the right footpad, and re-challenged them in the left footpad 16 days later, when the primary response was fully contracted (Figure 7E and Figure S9). The secondary challenge induced a massive re-expansion of the OT-II cells in the left dLN of WT mice. In contrast, the re-expansion was minimal when CD301b^+^ DCs were depleted immediately before the secondary immunization, indicating the requirement of CD301b^+^ DCs for the secondary expansion of the antigen-specific CD4T cells in the dLN (Figure 7F and 7G). Unlike the above experiments in which OT-II cells were adoptively transferred 1 day after the immunization, these experiments show the requirement of CD301b^+^ DCs for the optimal priming of preexisting clones since CD301b^+^ DCs were depleted only at the secondary immunization, eliminating the possibility that the abovementioned role of CD301b^+^ DCs is dependent on the timing of OT-II cell transfer. Collectively, these results indicate that CD301b^+^ DCs are required for optimal priming and expansion of rare antigen-specific CD4T cell clones.

## Discussion

The kinetics of antigen-specific CD4T cell priming is a critical determinant of immunological outcome in both quantitative and qualitative aspects of the response. For example, the frequency of antigen-specific CD4T cells in naive animals generally correlates with the response magnitude and reflects their repertoire diversity, which is tightly associated with the differentiation propensity through a TCR-intrinsic mechanism (Moon et al., 2007; Tubo et al., 2013). In addition, even among CD4T cells with fixed TCR specificity (i.e., TCR transgenic CD4T cells), the population size and the timing of priming significantly affect their participation in the memory response, indicating the number of antigen-specific CD4T cells initially recruited to the immune response as an independent parameter that influences the immunological outcome (Catron et al., 2006; Polonsky et al., 2018). However, while the molecular mechanism of CD4T cell trafficking in and out of LNs has been extensively studied, the cellular mechanism that maintains the number of CD4T cells in the reactive dLN during the priming is not well understood. By using a model for immunization with soluble protein antigens, here we show that CD301b^+^ DCs (1) facilitate the accumulation of lymphocytes in the dLN during the priming phase, (2) retain naïve CD4T cells through the MHCII-dependent interaction regardless of the TCR specificity, (3) directly present antigens to CD4T cells to induce early activation and cell cycle entry of antigen-specific CD4T cells, (4) promote Th2 differentiation while suppressing the Th1 and Th17 commitment, and (5) are required for the robust and timely expansion of rare antigen-specific CD4T cell clones during the primary and recall responses. These results reveal the critical importance of migratory CD301b^+^ DCs in CD4T cell priming dynamics in its early stage.

Immunization or infection in peripheral organs generally induces transient hypertrophy of the dLN. While DCs and CD4T cells cooperatively accelerate the cellular influx to the dLN and maximize the efficacy of the immune response (Kumamoto et al., 2011; Moussion and Girard, 2011; Webster et al., 2006), how the number of CD4T cells in the dLN is maintained during this process is less clear. Our data show that, while the donor B cells accumulate in the dLN nearly proportionally to the time between 2 and 72 hours after the transfer (26 and 96 hours post-immunization) into WT hosts, T cells accumulate far less efficiently compared with B cells, suggesting that the egress is a key mechanism for keeping the T cell numbers in the dLN in check in our model. Although it is generally assumed that immunization-induced dLN hypertrophy is associated not only with the increased cellular influx but also with the shutdown of lymphocyte egress from the dLN (Girard et al., 2012; Matloubian et al., 2004; Rivera et al., 2008; Schwab and Cyster, 2007; Shiow et al., 2006), earlier studies demonstrated that the egress shutdown, if any, is transient and is followed by a larger-than-usual efflux from the dLN, making the total flow volume (the number of lymphocytes that pass through the LN per time unit) much larger for the dLN than for the ndLN (Cahill et al., 1976; Hall and Morris, 1965). Nonetheless, despite the huge difference in total cellularity, our data show no major changes in LN dwell time of naive CD4T cells between the dLN and ndLN in WT hosts, suggesting the wide capacity range of LNs to scan incoming CD4T cells at a relatively constant rate. In fact, the egress of polyclonal CD4T cells from both dLN and ndLN in WT mice was non-linear, with no apparent loss of the input pool at least in the first 8 hours after the LN entry. Importantly, the CD301b^+^ DC-depleted mice lost the ability to hold CD4T cells during this 8-hour window in the dLN but not in the ndLN, suggesting that migratory CD301b^+^ DCs are mobilized to meet the increased demand for scanning polyclonal CD4T cells and function as an immunological “display window” that temporarily retains the incoming CD4T cells in the dLN. Importantly, this transient retention of polyclonal CD4T cells in the dLN is antigen-independent and is conceptually distinct from the well-characterized retention of T cells that occurs *after* their engagement with DCs presenting cognate peptides (Bousso, 2008). In accordance with our data, mathematical models also predict that the intensive scanning of T cells by DCs (thus longer LN dwell time) maximizes the priming efficacy when the antigen density and the frequency of antigen-specific clones are low (Lee et al., 2012). Of note, in contrast to the depletion of the total DC population that results in a severe hypotrophy of LNs due to a significant reduction in LN input (Kumamoto et al., 2011; Moussion and Girard, 2011), the depletion of CD301b^+^ DCs minimally affected the lymphocyte entry into the LNs.

To our surprise, the dLN transit time was similar between polyclonal CD4T cells and antigen-specific OT-II cells and was similarly reduced in CD301b^+^ DC-deleted mice as well as in CD301b^ΔMHCII^ mice. These data suggest that, in addition to the direct antigen presentation signals to the cognate CD4T cells, CD301b^+^ DCs provide MHCII-dependent, antigen-independent signals to induce a temporary delay in their dLN transit. In addition, in spite of the near complete CD69 upregulation regardless of the CD301b^+^ DC depletion, the majority of OT-II cells in our model left the dLN within 16 hours of their entry and before their initial cell division, indicating that, unlike the inflammation-induced egress shutdown (Shiow et al., 2006), the CD69 upregulation alone was not sufficient for preventing them from leaving the dLN. Previous studies have shown that continuous interaction of naive CD4T cells with DCs and self peptide-MHCII complexes in naive LNs slows down the LN transit of CD4T cells, induces partial expression of CD69 and helps naive CD4T cells to maintain their sensitivity to cognate peptides (Fischer et al., 2007; Hochweller et al., 2010; Mandl et al., 2012; Stefanova et al., 2002; Tomura et al., 2010). While such "sensitivity maintenance" of CD4T cells through the self peptide-MHCII interaction may also be provided by CD301b^+^ DCs, it is unclear if that alone can explain the delayed and impaired OT-II cell expansion in CD301b^+^ DC-depleted animals, since, unlike our study in which the priming status of OT-II cells was analyzed within a few hours after their transfer, the CD4T cells in these studies were typically pre-conditioned for 24 hours or longer in the MHCII- or DC-null environment. Further study is needed regarding the signaling events triggered upon antigen-independent engagement with CD301b^+^ DCs. Importantly, however, even under certain conditions in which LN-resident DCs might play a more predominant role than migratory DCs in presenting antigens to prime cognate CD4T cells (Gerner et al., 2017; Gerner et al., 2015), the antigen-independent role of CD301b^+^ DCs to retain polyclonal CD4T cells in the dLN may still be functional and relevant.

cDC2 cells, including CD301b^+^ DCs, are a group of heterogeneous, *Batf3*-independent cDCs that are often required for the differentiation of antigen-specific CD4T cells in to Th2 and/or Th17 effector cells (Gao et al., 2013; Kumamoto et al., 2013; Lewis et al., 2011; Murphy et al., 2016; Persson et al., 2013; Satpathy et al., 2013; Schlitzer et al., 2013; Tussiwand et al., 2015; Williams et al., 2013). cDC2 subsets activate antigen-specific CD4T cells more efficiently than the *Batf3*-dependent cDC1 cells, while cDC2 cells are less efficient in cross-presenting antigens to CD8T cells than cDC1 cells (Dudziak et al., 2007; Gerner et al., 2017; Vander Lugt et al., 2014). However, since cDC1 cells are required for Th1 differentiation under certain circumstances and have at least some capacity to present antigens to CD4T cells (Eickhoff et al., 2015; Hor et al., 2015; Igyarto et al., 2011; Martinez-Lopez et al., 2015), the relative contribution of cDC2 cell subsets to the CD4T cell priming kinetics has been unclear. Indeed, several studies have shown that the lack of either cDC subsets has little impact on the cell cycle progression of antigen-specific CD4T cells *in vivo* even when it significantly affects the differentiation of CD4T cells, while others showed a partial reduction in cell proliferation in the absence of a particular cDC subset (Igyarto et al., 2011; Kumamoto et al., 2013; Lewis et al., 2011; Martinez-Lopez et al., 2015; Vander Lugt et al., 2014). These studies suggest that, in stark contrast to the general requirement of cDC1 cells for CD8T cell priming (Edelson et al., 2010; Hildner et al., 2008), there is no single cDC subset that is absolutely required for CD4T cell priming. However, these previous studies typically preseeded the dLN with a large number of antigen-specific CD4T cells under continuous recirculation, making the interpretation of the cell division data much more complex compared with the data from the semi-synchronized priming presented in this study. Our data show that, while the antigen presentation to CD4T cells can be assumed by other antigen-presenting cell subsets, CD301b^+^ DCs play a unique role in efficient scanning of antigen specificity, timely priming and expansion, and Th2 differentiation of antigen-specific CD4T cells. Since their role appears to be partly dependent on their capacity to induce transient retention of polyclonal CD4T cells, potent inflammation could override their requirement if the inflammation-induced CD4T cell influx is massive enough. Nevertheless, under the type 1 immunization condition with CFA, CD301b^+^ DCs were still required for the maximal CD4T cell retention and expansion, but they also seem to have suppressed Th1 and Th17 differentiation. While the mechanism for such suppression remains to be elucidated, it may operate as a safety threshold to avoid overt activation of Th1- and/or Th17-driven inflammation and suggests potential cooperation between CD301b^+^ DCs and CD301b^−^ DCs in Th1/Th17 priming, which facilitate the retention of polyclonal CD4T cells and the induction of Th1/Th17 differentiation program, respectively. Given that the CD301b^+^ DCs represent a major subset of dermal-derived migratory DCs, it is reminiscent of the previously proposed model of the cooperative priming of CD4T cells by LN-resident and migratory DC subsets, though the latter model assumes the radio-resistant cells, such as epidermal Langerhans cells but not CD301b^+^ DCs, as the main migratory DC component (Allenspach et al., 2008).

Upon exposure to protease allergens or contact irritants, CD301b^+^ dermal DCs migrate to the dLN within 24 hours, a few days earlier than other skin-resident migratory DCs such as epidermal Langerhans cells and CD103^+^ dermal DCs (Allan et al., 2006; Kissenpfennig et al., 2005; Kumamoto et al., 2009; Kumamoto et al., 2013; Ochiai et al., 2014). This quick mobilization is facilitated by the direct sensing of the irritants by skin-innervating sensory neurons (Aderhold et al., 2020; Shiba et al., 2009; Shiba et al., 2012), but its immunological advantage has not been fully addressed. CD4T cell priming has been shown to be initiated by antigen-bearing migratory DCs at the outer edge of the T cell zone near the B cell follicles where the HEVs are densely distributed (Bajenoff et al., 2003; Baptista et al., 2019; Ingulli et al., 2002). Consistent with our observation, a recent study shows that the EBI2-dependent propensity of naive CD4T cells to localize to the outer T cell zone facilitates the priming of antigen-specific CD4T clones especially when the antigen dispersal is limited and/or when the cognate clones are rare (Baptista et al., 2019). Alternatively, they can be primed in the areas adjacent to the lymphatic sinuses by LN-resident cDC2 cells that sample antigens directly from the sinus (Gerner et al., 2017; Gerner et al., 2015). While these two modes of CD4T cell priming are not mutually exclusive, the latter mode does not require migration of DCs from the immunization site and is generally assumed to be responsible for the fast response, particularly when the antigen rapidly disperses through the lymphatics into the dLN that is preseeded by a sufficient number of antigen-specific CD4T cells. In contrast, the former migratory DC-dependent mechanism may be more important for antigens that are strictly restricted to the peripheral organs (Gerner et al., 2017; Gerner et al., 2015; Itano et al., 2003; Lee et al., 2009). Nevertheless, consistent with our observations, the full CD4T cell response often requires migratory DCs even when the CD4T cell priming by LN-resident DCs precedes the arrival of migratory DCs in the LN (Gerner et al., 2015; Itano et al., 2003).

In skin-dLNs, CD301b^+^ DCs account for a major subset of the dermal-derived migratory cDC2s (Kumamoto et al., 2009; Kumamoto et al., 2013). CD301b^+^ DCs are preferentially localized to near HEVs at the T-B border areas and are segregated from the epidermal-derived Langerhans cells and cDC1 cells that are localized in the central T cell zone, as well as from the LN-resident cDC2 cells that are associated with the lymphatic sinuses (Gerner et al., 2012; Gerner et al., 2015; Kissenpfennig et al., 2005; Kumamoto et al., 2009; Kumamoto et al., 2013; Stoltzfus et al., 2020). Thus, CD301b^+^ DCs in the skin-dLNs likely represent a subset of migratory DCs that interact with T cells on their entry route through the HEVs into the paracortex (Bajenoff et al., 2003). Notably, Th1 and Th2 cells in the LNs have been shown to be localized to the central area and the outer edge of the T cell zone, respectively (Leon et al., 2012; Liu et al., 2007; Randolph et al., 1999). While it is unclear whether Th1 and Th2 cells in these models arise from different clones that are initially primed by distinct DC subsets, we hypothesize, based on our data, that naive antigen-specific CD4T cells newly homed to the LN initially interact with CD301b^+^ DCs regardless of their later fate, and that those that additionally interact with cDC1 cells will acquire the Th1 cell fate, while those that remain in the interaction with CD301b^+^ DCs preferentially become Th2 cells. This sequential priming model for Th1 differentiation is also supported by the fact that Th1 differentiation is promoted by their expression of chemokine receptors CCR7 and CXCR3, both of which has ligands that are enriched in the central T cell zone and sinusoidal areas in the LNs and promote Th1 differentiation (Groom et al., 2012; Leon and Lund, 2019; Lian and Luster, 2015; Marsland et al., 2005; Randolph et al., 1999). While the Th1 differentiation may not require CD301b^+^ DCs *per se*, it is still supported by the extensive repertoire scanning by CD301b^+^ DCs. Taken together, our results demonstrate that the interaction with CD301b^+^ DCs dictates the dynamics of the CD4T cell response against soluble antigens in the LN. Further studies are needed to decipher their role in responses against particulate, cell-associated, or replicating antigens.

---

**Figure S1.**
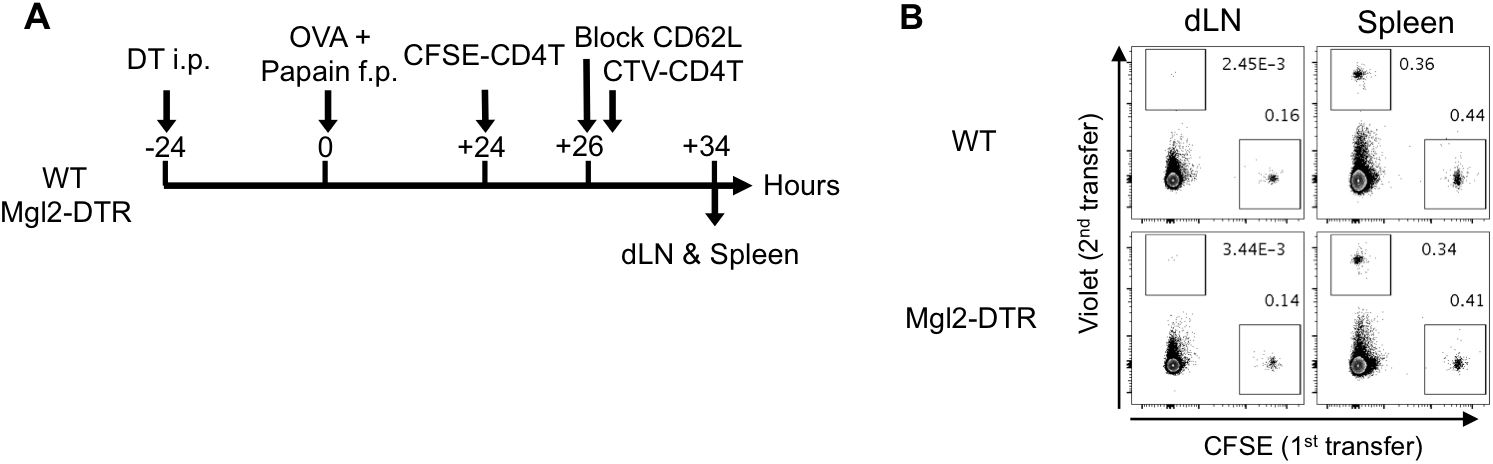
Related to Figure 2. Blockade of Lymphocyte Entry into LNs with Anti-CD62L mAb. (A, B) Total CD4T cells (2 × 10^6^) isolated from naive WT mice were labeled with CFSE and transferred into DT-treated Mgl2-DTR or WT mice that had been immunized with OVA and papain 24 hours earlier. The donor cells were allowed for 2 hours to home to LNs, after which further homing was blocked by retro-orbitally injecting anti-CD62L mAb. Immediately after the anti-CD62L injection, Cell Tracer Violet (CTV)-labeled CD4T cells (2 × 10^6^) were transferred and the dLN and spleen were harvested 8 hours after the CD62L blockade (A). Flow cytometry plots of CFSE+ and CTV+ CD4T cells in the dLN and spleen 8 hours after the CD62L blockade are shown (B).

**Figure S2.**
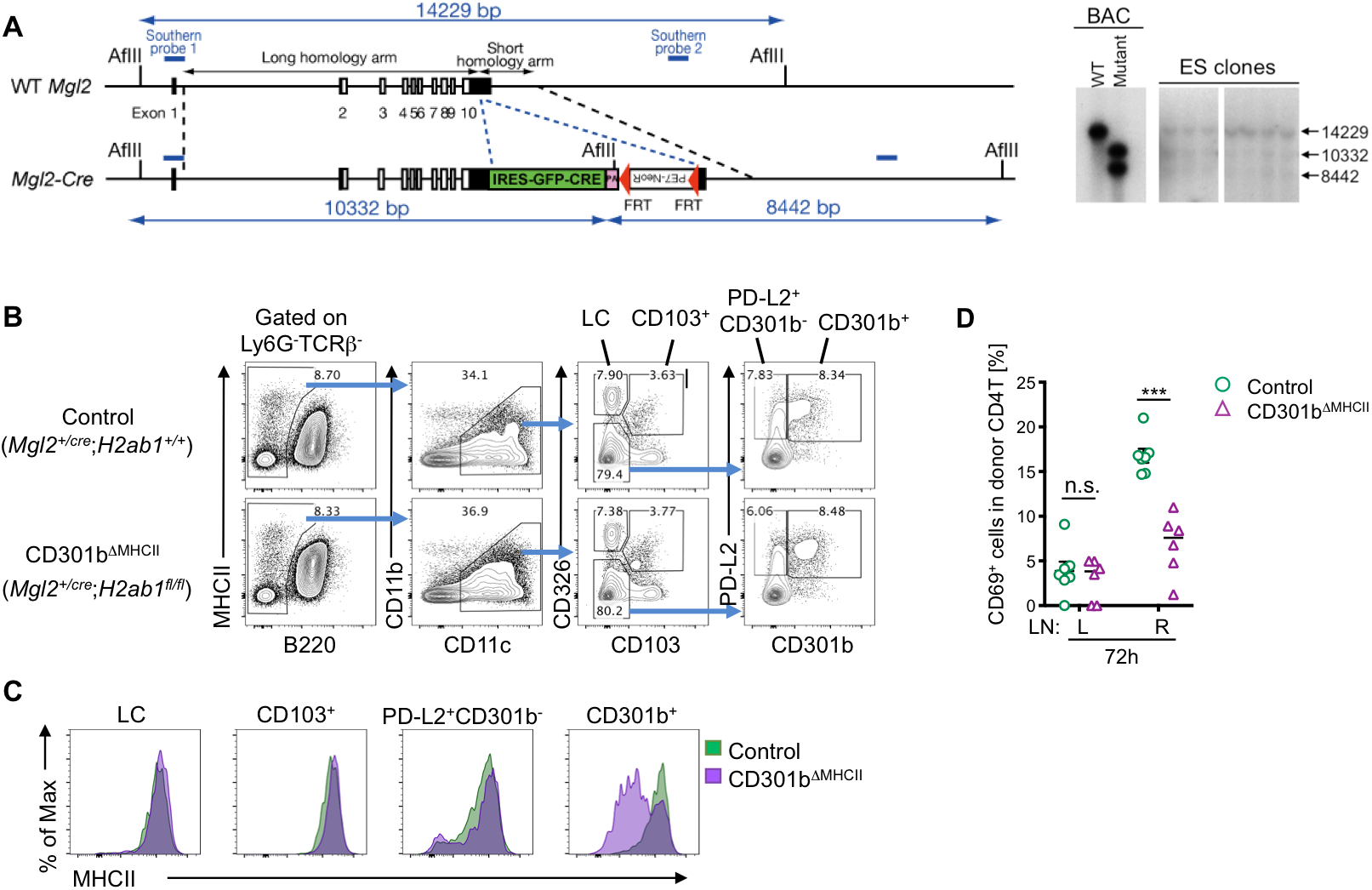
Related to Figure 2. Generation of the CD301b^ΔMHCII^ Mouse. (A) Targeting strategy for generating the Mgl2-Cre mouse. The targeting construct was created by editing the bacterial artificial chromosome (BAC) RP23-67I17 and retrieved onto the targeting vector. Successfully recombined ES cell clones were selected by Southern blotting. (B, C) CD301b^+^ DC-specific deletion of MHCII in the CD301b^ΔMHCII^ mice. Skin-dLNs were harvested from naive CD301b^ΔMHCII^ or control mice and stained for DC subsets as shown in (B). Representative flow cytometry histograms showing MHCII expression in each DC subset are shown in (C). MHCII was specifically deleted in CD301b^+^ DC population but remained intact in the other DC subsets including Langerhans cells (LC), CD103^+^ DCs, and PD-L2^+^ CD301b^−^ DCs. (D) As in Figure 1, WT splenocytes (1 ×10^7^ cells) were transferred into CD301b^ΔMHCII^ mice that had been immunized with OVA and papain in the right hind footpad 24 hours earlier. Right draining (R) and left non-draining (L) popliteal LNs were harvested 72 hours after the transfer. CD69 expression in the donor cells is shown.

**Figure S3.**
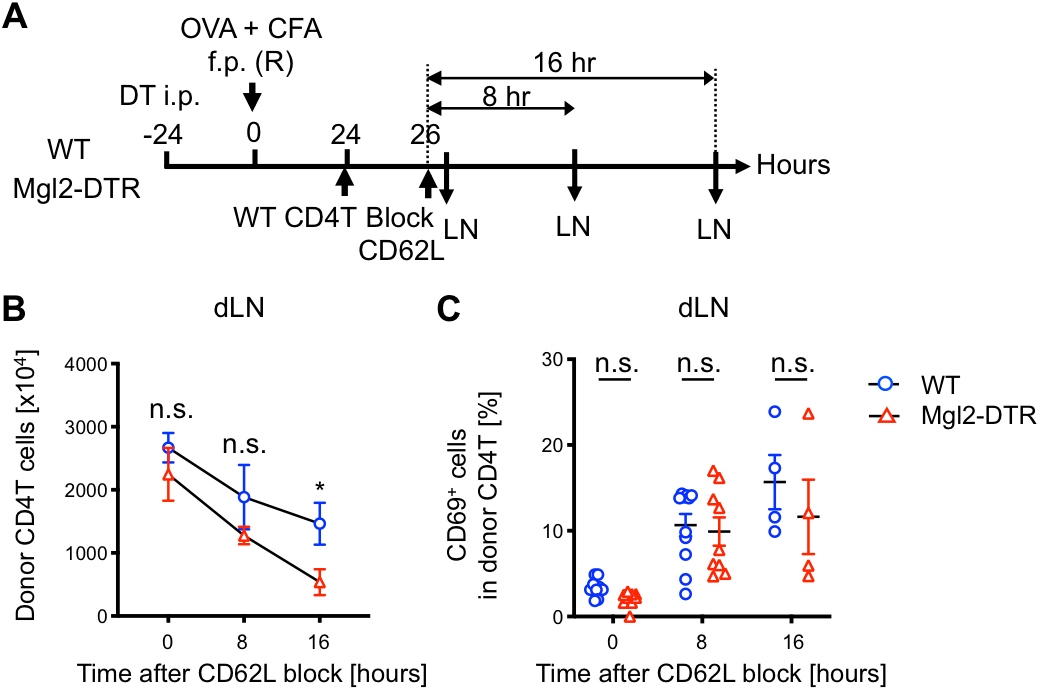
Related to Figure 2. CD301b^+^ DCs Retain Naive Polyclonal CD4T Cells in the dLN under Non-Th2 Immunization Condition. (A) As in Figure 2A, but the donor mice were immunized with OVA and CFA. (B, C) Number (B) and CD69 expression (C) of the donor CD4T cells in the dLN at indicated time-points. Data are pooled from 3–6 mice per group at each time point. Data represent mean ± SEM. *p < 0.05, n.s. not significant, by two-tailed Student’s t test.

**Figure S4.**
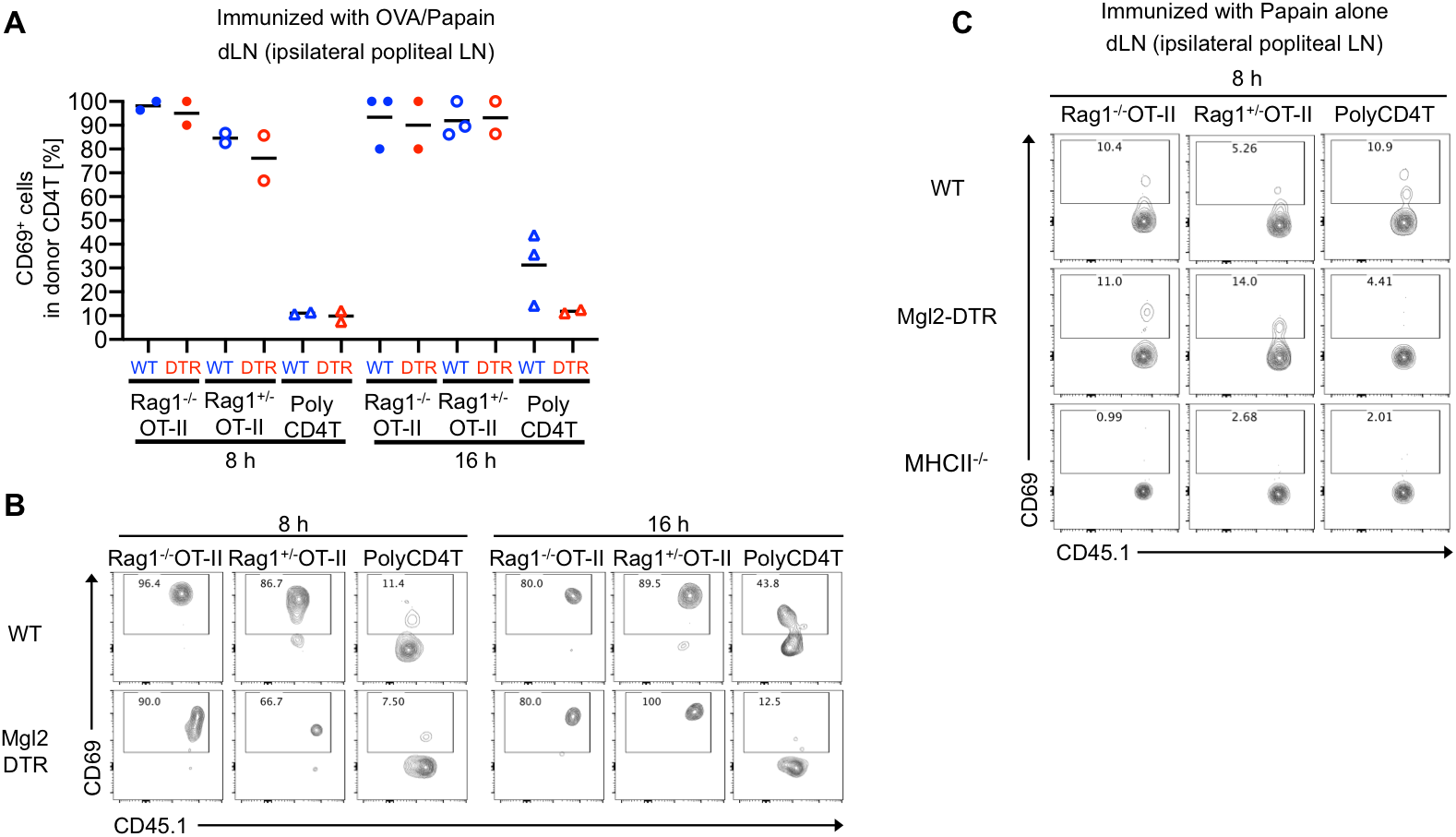
Related to Figure 3. MHCII- and antigen-dependency of CD69 upregulation in *Rag1*^*−/−*^ OT-II, *Rag1*^*+/−*^ OT-II, and polyclonal CD4T cells. (A-C) As in Figure 3, but *Rag1*^*−/−*^ OT-II, *Rag1*^*+/−*^ OT-II, and WT polyclonal CD4T cells (1 × 10^6^ cells each) were co-transferred into DT-treated Mgl2-DTR or WT recipients that had been immunized as indicated. In (C), the donor cells were also transferred into a group of MHCII^*+/−*^ recipients.

**Figure S5.**
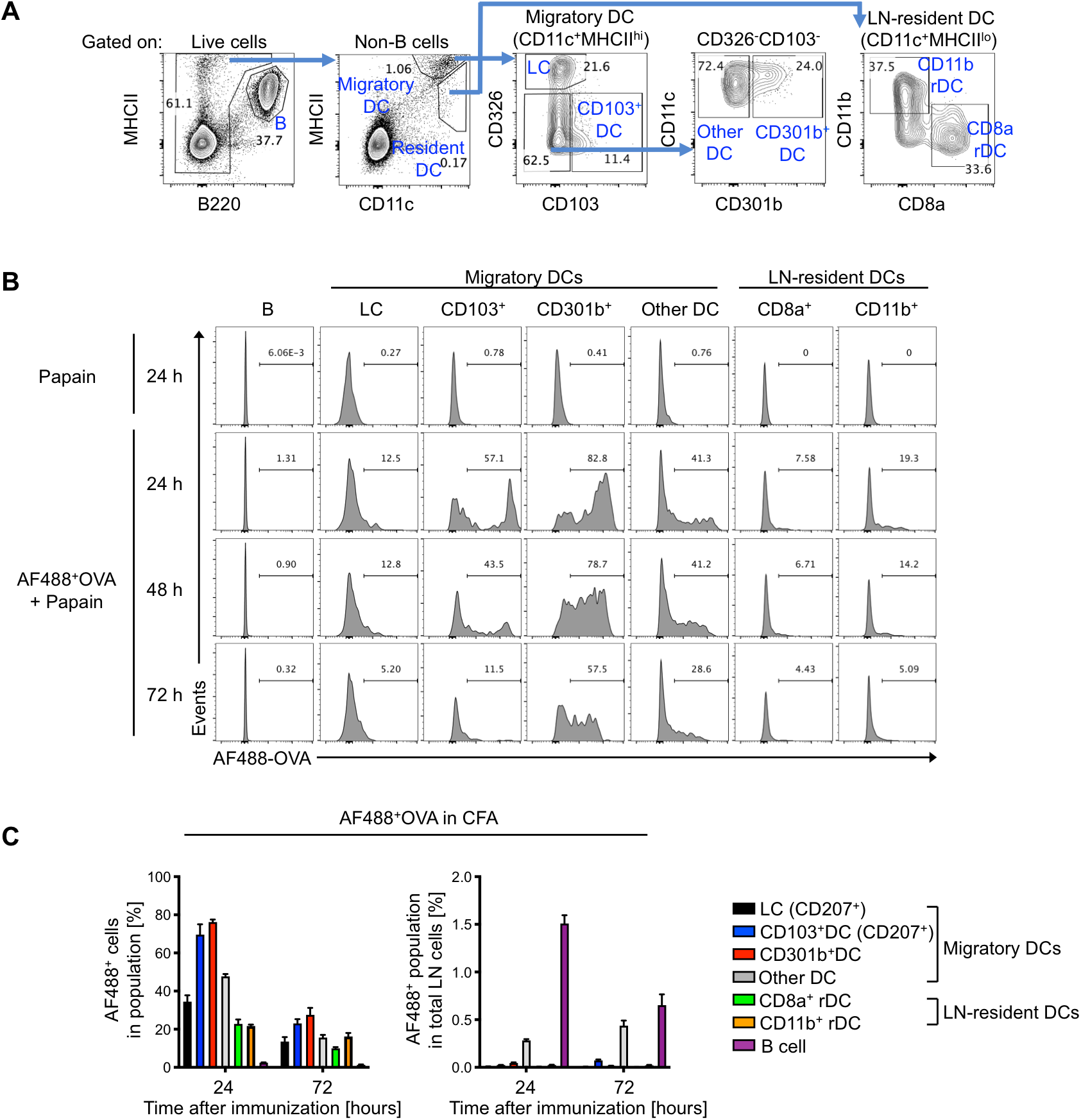
Related to Figure 4. Antigen Uptake by CD301b^+^ DCs in the dLN Early After Immunization. (A-C) WT mice were immunized with AF488-labeled OVA protein with papain (A, B) or with CFA (C) in the footpad and the dLNs were harvested and stained for DC subset markers at indicated time-points. (A) Gating strategy used to identify DC subsets. (B) Representative flow cytometry histograms for AF488-OVA uptake by each cell type identified in (A). (C) Frequency of AF488^+^ DC subsets in the dLN at indicated time-points after immunization with OVA and CFA.

**Figure S6.**
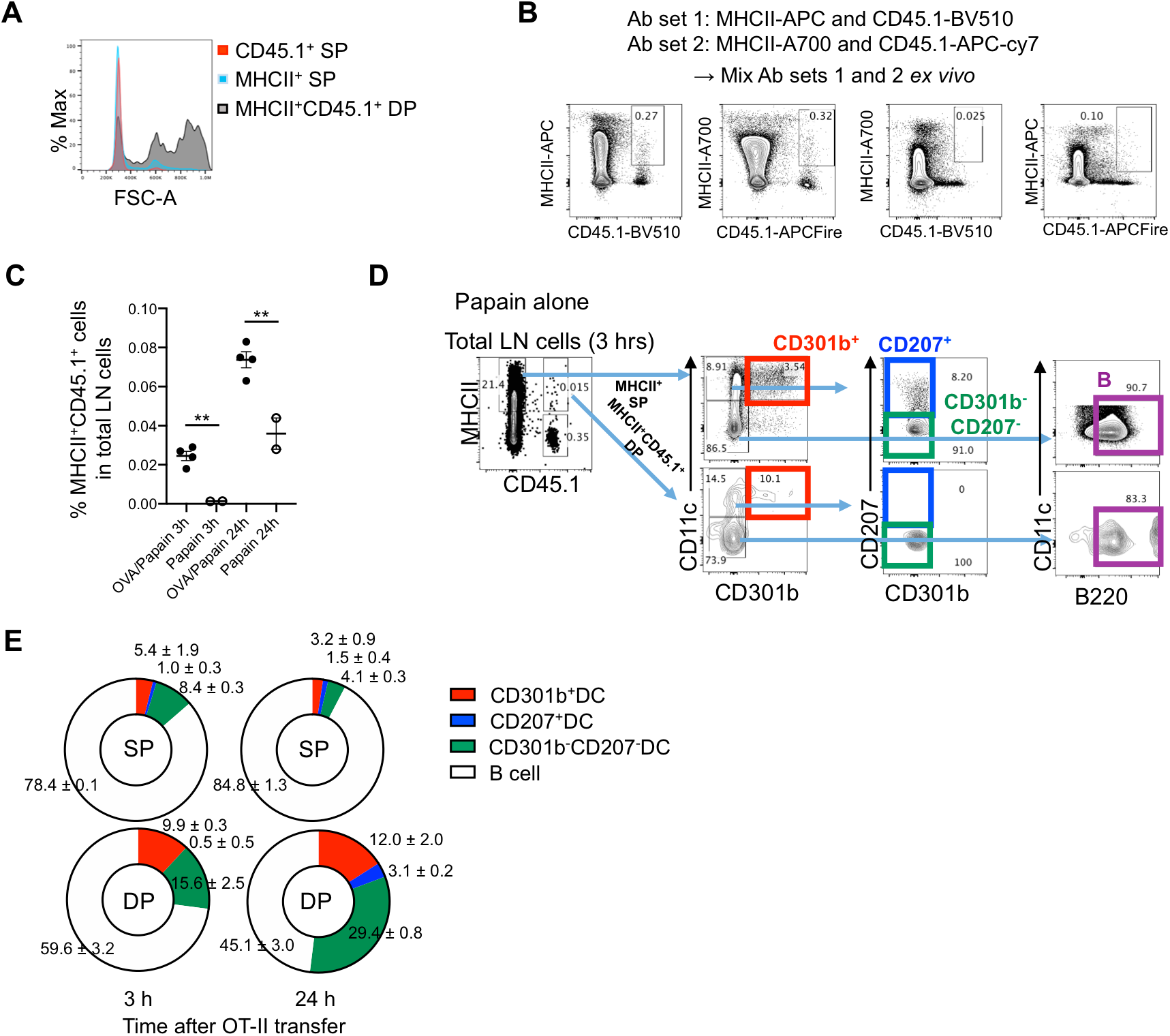
Related to Figure 4. Direct Antigen Presentation by CD301b^+^ DCs to CD4T cells in the dLN Early After Immunization. (A, B) Mice were immunized as in Figure 4F. Forward scatter (FSC-A) of the CD45.1^−^ MHCII^+^ single-positive (SP) or CD45.1^+^ MHCII^+^ double-positive (DP) events in the dLN cells 3 hours after the OT-II cell transfer (A). In (B), two sets of LN cells were separately stained for CD45.1 and MHCII with a different set of fluorochromes (MHCII-APC and CD45.1-APC-Cy7, or MHCII-A700 and CD45.1-BV510). After washing, the two sets of cells were mixed together and incubated for another 10 min. Flow cytometric plots of MHCII and CD45.1 staining are shown. Note that fluorochrome exchange between the two sets of cells is minimal. (C-E) As in Figure 4F but the mice were immunized with papain alone. The frequency of the total CD45.1^+^ MHCII^+^ DP events was compared to those immunized with OVA and papain (C). In (E), the composition of MHCII^+^ cell subsets among the CD45.1^−^ MHCII^+^ SP or CD45.1^+^ MHCII^+^ DP events was calculated based on the gating strategy shown in (D). Data indicate mean ± SEM (C, E) or representative flow cytometry plots of at least two independent experiments (A, B, D). **p < 0.01 by two-tailed Student’s t test.

**Figure S7.**
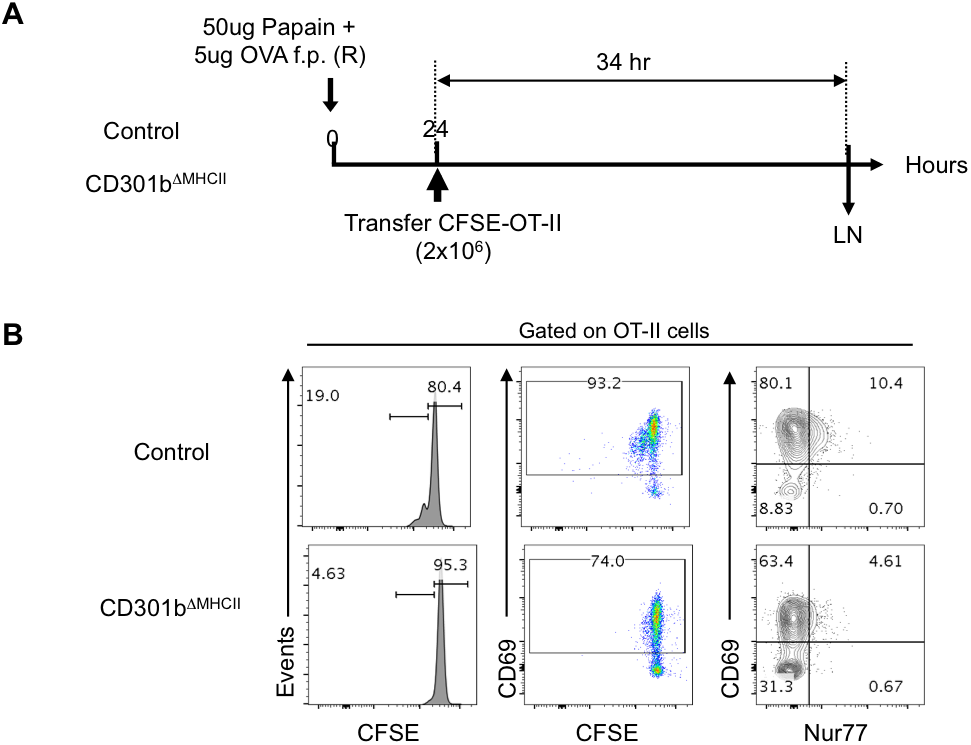
Related to Figure 5. Requirement of MHCII expression by CD301b^+^ DCs for Optimal Priming of Antigen-Specific CD4T Cells. CD301b^ΔMHCII^ or control mice were immunized with OVA and papain in the footpad and transferred with CFSE-labelled OT-II cells 24 hours post immunization. The dLNs were harvested 34 hours after the transfer and stained for the cell surface expression of CD69 and intracellular expression of Nur77. (A) Experimental design. (B) CFSE dilution (left), CD69 expression (middle) and Nur77 expression (right) in the OT-II cells. Cell cycle entry, CD69 upregulation and Nur77 expression by the donor OT-II cells were significantly delayed and diminished in CD301b^ΔMHCII^ mice.

**Figure S8.**
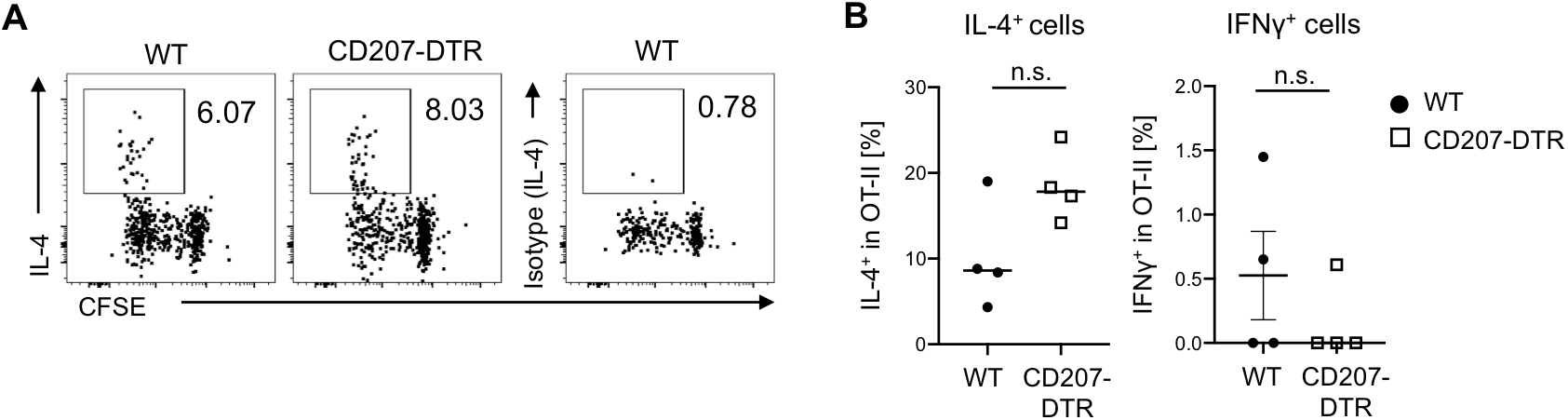
Related to Figure 6. CD207^+^ DCs are Dispensable for the Th2 Differentiation of Antigen-specific CD4T Cells upon Immunization with OVA and Papain. (A, B) DT-treated WT or CD207-DTR mice were transferred with 1 × 10^5^ CFSE-labeled OT-II cells and immunized with OVA and papain in the footpad. The dLNs were harvested 7 days after the immunization and restimulated *in vitro* with PMA and ionomycin for intracellularly staining IL-4 and IFNγ. Data represent mean ± SEM (B) or show representative flow cytometry plot of at least two independent experiments (A). n.s. not significant by two-sided Student’s t test.

**Figure S9.**
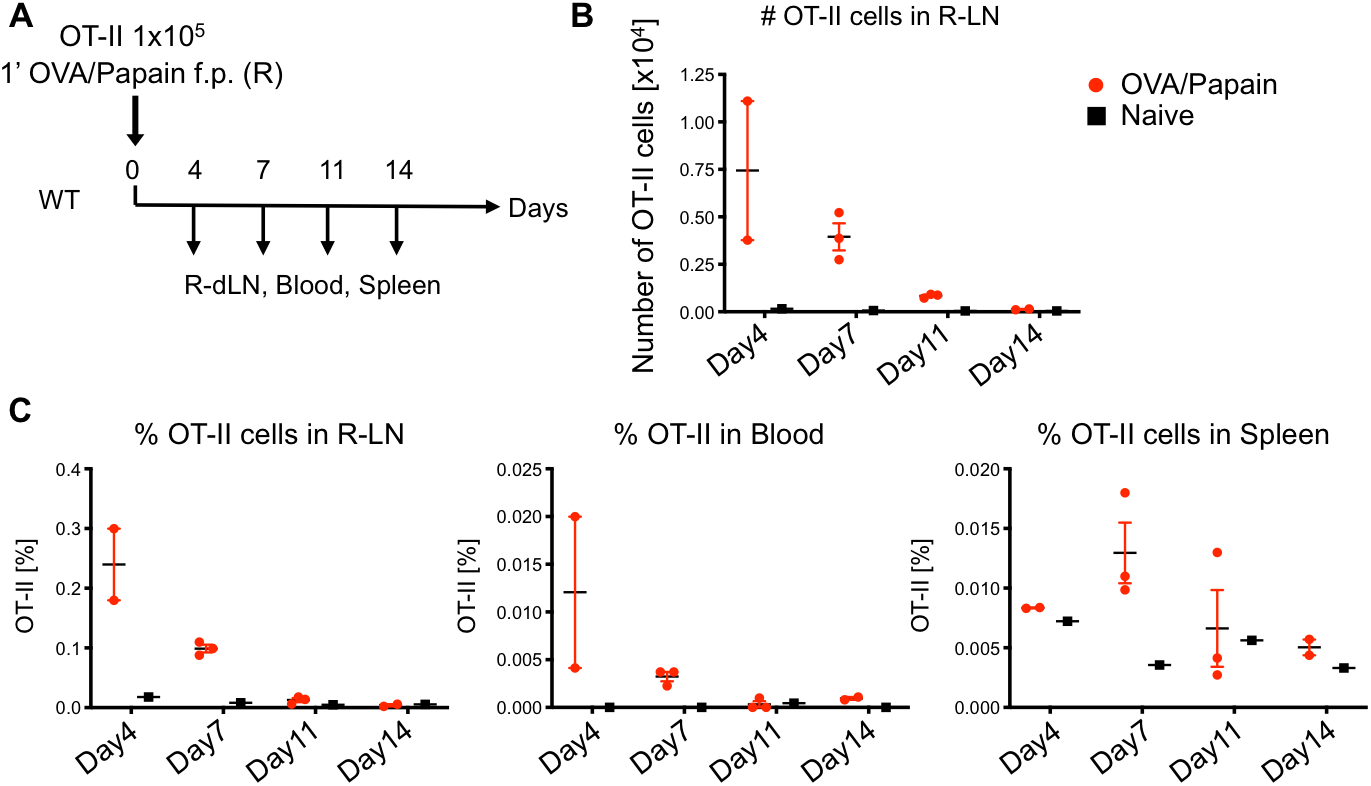
Related to Figure 7. Kinetics of OT-II Cell Expansion and Contraction Following Primary Immunization with OVA and Papain. WT or Mgl2-DTR mice were transferred with 1 × 10^5^ OT-II cells and immunized with OVA and papain in the right footpad or left unimmunized (Naive). OT-II cell numbers in the dLNs, blood, and spleen were examined at indicated time-points as in (A). The number (B) and percentage (C) of OT-II cells in the dLNs, blood and spleen at indicated time-points are shown. Data represent mean ± SEM.

## Acknowledgements

We thank Akiko Iwasaki for Mgl2-Cre and Mgl2-DTR mice. We also thank William Gause and George Yap for critical reading of the manuscript and Jihad El-Fenej and Alejandro Davila-Pagan for technical assistance. This work was supported by NIH grant AI132576 (Y.K.).

## Author Contributions

N.T. and Y.K. designed experiments. N.T., A.L.C. and Y.K. conducted experiments. N.T. and Y.K. analyzed and interpreted the data. Y.K. conceptualized and supervised the study. N.T. and Y.K. wrote the paper.

## MATERIALS AND METHODS

### Mice

Mgl2-DTR and *Mgl2*^*Cre*^ mice were a gift from Akiko Iwasaki (Yale University). Generation of *Mgl2*^*Cre*^ mice is described below. C57BL/6 (B6) and congenic CD45.1 (B6.SJL-PtprcaPep3b/BoyJ) mice on B6 background were purchased from Charles River Laboratory and propagated in our colony. MHCII^fl/fl^ (B6.129X1-H2-Ab1^tmKoni^/J), MHCII^−/−^ (B6.129S2-H2^dlAb1-Ea^/J), OT-II (B6.Cg-Tg(TcraTcrb)425Cbn/J), CD207-DTR (B6.129S2-Cd207^tm3(DTR/GFP)Mal^/J), *Rag1*^*−/−*^ (B6.129S7-Rag1tm1Mom/J) and Nur77-GFP reporter (C57BL/6-Tg(Nr4a1-EGFP/cre)820Khog/J) mice were purchased from the Jackson Laboratory (Bar Harbor, ME) and maintained in house. *IFNAR*^*−/−*^ mice were a gift from Karen Edelblum (Rutgers New Jersey Medical School). Mgl2-DTR mice (B6.*Mgl2^+/DTR-EGFP^*) were previously described and maintained in our colony (Kumamoto et al., 2013). OT-II mice were crossed with CD45.1 mice to generate CD45.1;OT-II mice. The CD45.1;OT-II mice were maintained on the *Rag1*^*−/−*^ background and used for some experiments. Nur77-GFP mice were crossed with CD45.1 or CD45.1;OT-II mice to generate CD45.1;Nur77-GFP or CD45.1;Nur77-GFP;OT-II mice, respectively. All animal experiments in this study have been approved by the Institutional Animal Care and Use Committee at Rutgers New Jersey Medical School.

### Generation of CD301b^ΔMHCII^ mice

A targeting construct was designed to introduce an IRES-EGFP-Cre cassette in the 3’ untranslated region of the *Mgl2* gene. The IRES-EGFP-Cre construct was cloned from the pIGCN21 plasmid (gift from Neal Copeland) (Lee et al., 2001). To generate the *Mgl2*^*Cre*^ mice, an IRES-EGFP-Cre-polyA cassette linked to a neomycin-resistant cassette (NeoR) flanked by FRT sites was targeted into the 3’ untranslated region of the *Mgl2* gene, downstream of the stop codon in the last exon (exon 10). To that end, the IRES-EGFP-Cre-polyA cassette was first cloned into a bacterial artificial chromosome DNA containing the *Mgl2* allele (R23-67I17) and then retrieved onto a targeting vector as previously described (Kumamoto et al., 2013). C57BL/6N embryonic stem (ES) cells were electroporated with the linearized targeting vector. After selection in G418, the ES clones were screened for proper recombination by Southern blot with previously described probes (Kumamoto et al., 2013), and microinjected into blastocysts of albino B6 mice. The germline-transmitted offspring of the chimeric mice was once crossed wtih mice expressing an FLP recombinase (B6.129S4-Gt(ROSA)26Sortm2(FLP*)Sor/J, Jackson Laboratories) to remove the NeoR cassette. The resulting *Mgl2*^*+/Cre*^ mice were crossed together to generate homozygous *Mgl2^cre/cre^* mice, which were then crossed with MHCII^fl/fl^ mice to generate *Mgl2*^*+/Cre*^;MHCII^fl/fl^ (CD301b^ΔMHCII^) mice. We noted a few CD301b^ΔMHCII^ mice had global deletion of MHCII in the whole body due likely to the sporadic low-level expression of the Cre in germ cells or during early embryogenesis as reported for other Cre/loxP systems (Heffner et al., 2012; Liu et al., 2019). While the events were rare, the blood B cells of all CD301b^ΔMHCII^ mice and the control mice were examined by flow cytometry to remove any mice with the global MHCII deletion from the analysis. Global deletion of the *H2-Ab1* gene was further examined by tail DNA genotyping using a mixture of forward (5’- CTCTACACCCCCAACACACC-3’) and two different reverse (5’- TCGCCTTCTTGACGAGTTCT-3’ and 5’- ACTCTCTGGTCTCCGAACGA-3’) primers, allowing amplification of a 200-bp band and a 140-bp band corresponding to the intact and recombined *H2-Ab1* alleles, respectively.

### DT treatment and immunization

For DC depletion, mice were injected with 500 ng DT (List Biological Laboratories) intraperitoneally at indicated time-points. All immunization procedures were performed in the rear footpad with 20 μL injection volume per footpad as described previously (Kumamoto et al., 2013). Mice were immunized as indicated in each Figure with 5 μg low-endotoxin OVA (Worthington Biochemical Corporation) together with 50 μg papain (P4762, Sigma) or 10 μL CFA (F5881, Sigma). For *in vivo* antigen-uptake experiments, OVA was labeled with Alexa Fluor 488 Protein Labeling Kit (A10235, Thermo Fisher) and injected (5 μg) in the rear foot pad together with 50 μg papain or with 10 μL CFA. The ipsilateral (dLN) and contralateral (ndLN) popliteal LNs were harvested at indicated time-points.

### Cell trafficking and priming analysis

For all cell transfer experiments, the donor cells were isolated from naive mice and the indicated number of cells were transferred retro-orbitally in 0.5 mL suspension in PBS under light anesthesia with isoflurane.

For splenocyte accumulation assay, donor splenocytes were isolated from naive B6 or *IFNAR*^*−/−*^ mice by digesting the spleen with 2.5 mg/mL collagenase D (11088882001, Sigma) at 37°C for 30 min and labeled with 1.0 μM CFSE (eBioscience 65-0580-84, Thermo Fisher) according to the manufacturer’s protocol. Ten million cells were transferred into recipient mice that had been treated with DT and immunized as indicated. The left (ndLN) and right (dLN) popliteal LNs were harvested 2 or 72 hours after the transfer.

For analyzing the CD4T cell priming kinetics, CD4T cells were negatively isolated from the digested spleen and LNs of indicated donor mice with Mouse CD4T Cell Isolation Kit (STEMCELL Technologies 19852 or BioLegend 480033) according to the manufacturer’s protocol and labeled with 1.0 μM CFSE (Thermo Fisher) or CellTrace Violet (C34571, Thermo Fisher). The purity was typically 90-95%. Two million cells were transferred into recipient mice that had been immunized 24 hours prior and treated with DT as indicated. In some experiments, multiple types of donor cells (1 × 10^6^ cells each) were co-transferred into one mouse. Two hours after the transfer, further LN entry of T cells was blocked by retro-orbitally injecting 200 μg anti-CD62L mAb (clone Mel-14, BE0021, BioXCell). The left (ndLN) and right (dLN) popliteal LNs were harvested at indicated time-points.

For analyzing the OT-II cell priming in CD207^+^ DC-depleted mice, CFSE-labeled OT-II cells (1 × 10^5^) were adoptively transferred into DT-treated WT or CD207-DTR mice, followed by immunization with OVA and papain in the footpad. The dLNs were harvested on day 7 post-immunization. For analyzing the OT-II cell priming in CD301b^ΔMHCII^ mice, CFSE-labeled OT-II cells (2 × 10^6^) were adoptively transferred into CD301b^ΔMHCII^ or control *Mgl2*^*+/Cre*^;*H2ab1^+/+^* mice that had been immunized with OVA and papain for 24 hours. The dLNs were harvested for analysis 34 hours after the transfer. For analyzing the priming efficiency of OT-II cells, titrated numbers of OT-II cells (1, 10, 100, 1,000, or 10,000) were mixed with naive polyclonal CD4T cells so that the total donor cell number is 10,000 and transferred into the indicated recipients that had been immunized 1 day prior. The dLNs were harvested 4 days after the transfer. Some mice were treated with DT on days −1 and +2.

For analysis of OT-II cell re-expansion in response to secondary immunization, WT or Mgl2-DTR mice were transferred with OT-II cells (1 × 10^5^) and immunized with OVA and papain in the right footpad on day 0. The mice were then treated with DT on day 16, followed by a secondary challenge with OVA and papain in the left footpad on day 17. The right and left popliteal LNs were harvested 3 days after the secondary challenge. Contraction of the primary response was verified before the secondary immunization by counting the number of circulating OT-II cells in a separate group of WT mice that had been immunized side-by-side with the experimental group.

### Cell preparations and flow cytometry

For staining cell surface antigens, LNs were minced and digested with 2.5 mg/ml collagenase D in complete DMEM with 10% FBS at 37°C for 30 min. In the case of spleen, erythrocytes were lysed by briefly suspending cells in ACK lysis buffer. Cells were resuspended in 2 mM EDTA in PBS and stained with cell viability dye (Zombie Aqua or Zombie UV, BioLegend). Cells were then incubated with 10 μg/mL anti-CD16/CD32 (2.4G2, BioLegend) on ice for 10 min to block non-specific antibody binding to Fc receptors and stained with fluorochrome-conjugated monoclonal antibodies (mAbs) on ice for 20 min. All staining mAbs were prepared in FACS buffer (1% BSA, 2mM EDTA, 0.05% sodium azide in PBS). For detecting T cell-DC conjugates, the minced LNs were gently mushed with a plunger without collagenase digestion and resuspended in PBS without EDTA.

For intracellular cytokine staining, cells were stimulated in a 96-well round-bottom plate with Cell stimulation cocktail containing PMA and ionomycin (eBioscience 00-4970-03, Thermo Fisher) at 37°C for 1 hour, then incubated another 5 hr at 37 C with additional Protein Transport Inhibitor Cocktail (eBioscience 00-4980-03, Thermo Fisher). Cells were then fixed and permeabilized with BD Cytofix/Cytoperm Kit (BD Biosciences) and incubated with anti-cytokine mAbs for 30 min on ice. For staining Nur77 by mAb, cells were fixed and permeabilized with Foxp3 Fixation/Permeabilization Concentrate and Diluent (eBioscience 00-5521-00, Thermo Fisher).

The following antibodies were used for flowcytometry analysis: Anti-CD4 (RM4-5 or GK1.5), CD8a (53-6.7), CD45R/B220 (RA3-6B2), TCRβ (H57-597), Ly6G (1A8), CD69 (H1.2F3), MHCII (M5/114.15.2), CD11b (M1/70), CD11c (N418), CD326 (G8.8), CD301b (URA1), CD207 (4C7), CD45.1 (A20), CD45.2 (104), Nur77 (12.14), IFNγ (XMG1.2), IL-17A (TC11-18H10.1), IL-4 (11B11) and CD124 (I015F8) mAbs were purchased from BioLegend. Anti-CD44 (IM-7), CD301b (11A10-B7), CD103 (2E7), and PD-L2 (TY25) mAbs were purchased from eBioscience. Flow cytometry was performed on BD LSRII (BD Biosciences) or Attune NxT (Thermo Fisher) and analyzed by FlowJo software (Version 9.3.2 and 10.5.0, BD).

### Immunohistochemistry

Axillary and brachial lymph nodes were harvested from WT mice one day after immunization with OVA and papain in the front footpad and frozen in OCT compound (Sakura Finetek). Cryosections 5-7 μm in thickness were cut and fixed in ice-cold acetone, and non-specific binding was blocked by pre-incubating the sections with 2 % normal goat serum in PBS. For staining DC subsets and HEV, fixed sections were first incubated with 5 μg/ml anti-CD301b (clone URA1, Invitrogen), anti-CD207 (clone eBioRMUL.2, Invitrogen), or biotin-conjugated anti-CD11c (clone N418, BioLegend) mAbs. HRP-conjugated goat anti-rat IgG γ chain-specific antibody (Jackson ImmunoResearch) was used to detect the anti-CD301b and anti-CD207 mAbs and HRP-conjugated streptavidin (SA-HRP, Biotium) was used to detect the biotinylated anti-CD11c mAb. The HRP-conjugated secondary antibody and streptavidin were detected by Alexa Fluor 488 Tyramide (Biotium). The sections were then incubated with 2.5 μg/ml anti-PNAd (clone MECA79, Biolegend), followed by HRP-conjugated anti-rat IgM μ chain-specific antibody (Jackson ImmunoResearch) and Alexa Fluor 647 Tyramide (Biotium). For double staining CD301b^+^ DCs and B cells, the sections stained for CD301b were incubated with 2.5 μg/ml biotinylated CD45R/B220 (clone RA3-6B2, BioLegend), followed by detection with SA-HRP and Alexa Fluor 647 Tyramide. The nuclei were stained with DAPI (BioLegend). The sections were mounted in Fluoromount G (Diagnostic BioSystems) and analyzed with a BZ-X710 fluorescence microscope (KEYENCE). For measuring the distance from DCs to the nearest HEV, the distance between the centroid of each DC and the outline of the nearest HEV was measured after segmentation of DCs and HEVs using ImageJ (version 2.1.0/1.53c) with a plugin DiAna (Gilles et al., 2017). All positions of DCs analyzed were confirmed to be outside of the segmented HEVs.

### Statistical Analysis

P-values were calculated by two-tailed Student’s t test by Prism software (version 7, GraphPad). Differences were defined as statistically significant when P-values were <0.05. Data are presented as mean ± SEM.

